# Novel Exploration Growth Quantifies Anxiety-like Behaviour in the Elevated Plus Maze

**DOI:** 10.1101/2024.06.10.598202

**Authors:** Matthew D. Zelko, Stephen R. Robinson, Elisa Hill-Yardin, Helen Nasser

## Abstract

The elevated plus maze is intended to provoke anxiety-like behaviour, which is primarily measured by the degree to which the rodent explores or avoids the risker, unenclosed arms. Measures such as arm entries and total time spent within the arm are conventionally used, but their analysis often produces inconsistent inferences about the level of anxiety-like behaviour being observed. This inconsistency occurs because the measures do not correlate with one another, raising the question of how validly they capture both the exploratory and avoidance motivations that typify anxiety-like behaviour. Given this inconsistency, we propose a new measure, novel exploration growth, that captures avoidance and exploratory behaviours within the maze. The growth of novel exploration tracks the first visit to discrete areas of the maze over time. The absolute amount of novel exploration, combined with the phasic nature of the exploration, reveals behavioural phenotypes with both avoidance and exploration quantified in a single time series. By addressing both motivations evoked by the maze through a single coherent measure, we provide a superior estimation of anxiety-like behaviour and resolve inconsistencies that arise when applying conventional measures alone.

## Introduction

The elevated plus maze (EPM) is a popular paradigm for testing rodents’ anxiety-like behaviour (ALB). The EPM consists of four elevated arms, two of which are surrounded by high walls, known as closed arms. The remaining two arms, known as the open arms, are not enclosed. These two distinct arm types allow researchers to observe whether the rodent seeks safety in the closed arms or risk detection by exploring the open arms. (Carola et al., 2002; Leo & Pamplona, 2014). Avoidance of the open arms in favour of the safety of the closed arms is considered to indicate a greater degree of ALB. In contrast, exploration of the open arms indicates a lower degree of ALB. Exposure to the maze, therefore, evokes a conflict between the innate motivations of safety and exploration, characterising the ALB as an indicator of emotional state (Hoffman et al., 2022; Perusini & Fanselow, 2015).

Several measures have been used to compare differences in preferencing of an arm type. The standard measures include the total number of entries and time spent in an arm type. As these measures lack temporal sensitivity, researchers also track the initial latency to enter an arm type (Walf & Frye, 2007). However, using latency to enter an arm type assumes that every entrance into an arm is comparable, ignoring differences in the depth of exploration that follows it. This limitation can result in an early, shallow entry into the open arms, which is interpreted as lower anxiety-like behaviour than a delayed but more deep exploratory entry (Tejada et al., 2009).

Alongside the lack of temporal or spatial sensitivity of these conventional measures, it is unclear how to prioritise outcomes from these measures when making inferences about the level of ALB being observed in rodents. For example, a rodent may only enter the open arms a few times but may spend more time investigating the environment in depth. Ethologically, this rodent displays high amounts of exploratory drive. Conversely, another rodent may make many shallow entries into the open arms but never fully commit to exploring them beyond the entry boundary, a pattern that should be inferred as a higher degree of ALB. Despite this, the greater number of entries into the open arms would typically be interpreted as a lower degree of ALB compared to the former exploratory counterpart. Thus, the most appropriate selection of available behavioural measures is contingent upon potentially subjective decisions determining which behaviour best typifies ALB. This need for subjectivity could lead to difficulty in replicating outcomes or translating intervention effects to human populations.

The present paper overcomes the abovementioned limitations by providing a single, coherent measure of rodent exploratory and avoidant behaviours in the EPM. Using a new measure of ALB, novel exploration growth (NEG), safety and exploration motivational preferences are observed and differentiated into qualitatively distinct behavioural phenotypes. By measuring NEG over time, each behavioural phenotype is mapped to an appropriate level of ALB to allow justified and replicable inferences to be made about the traversal behaviours of rats in the EPM.

A multistage analysis workflow enables inferences to be made about the level of ALB using NEG. This workflow includes collecting and preprocessing tracking data, which are then analysed using Bayesian change-point regression and Bayesian generalised additive model (GAM) regression. Change-point analysis is used to identify the phases of exploratory behaviour during testing. These phases are then analysed using Bayesian GAM regression to estimate the effects of interest (e.g., between phenotypes or arm types).

To demonstrate how to apply the proposed workflow, distinct behavioural phenotypes were simulated across the entire EPM and for each arm type and their NEGs were analysed. In addition, we compared the three phenotypes across the conventional measures to compare the difference in ALB level inferred from these measures and NEG. This comparison highlights the incongruence between these conventional measures and demonstrates how NEG, as a unified and coherent measure, remedies these inconsistencies.

### Novel Exploration Growth as a Unified Measure of Exploration and Avoidance

NEG is measured by tracking the first physical visit to discrete areas of the EPM. The arena is divided into a 1 cm × 1 cm grid in a similar manner to that used in previous studies (Costa & Tinós, 2016; Salum et al., 2000). The initial visit to each square is recorded, and unique visits are aggregated over time and expressed as a percentage of the total grid units of the arm type of interest or the entire maze. For example, if a rodent has physically visited all of the available grid squares in the closed arms and half of the available squares in the open arms at a given point in time, they would have a total NEG of 75%, a closed arm NEG of 100% and an open arm NEG of 50%.

The choice to use the first visit to a grid square is based on the utility of the existing latency to enter the arm measure to the entire arm or maze. This utility exists because there is a need to distinguish between naïve exploration and informed preference when evaluating the motivation to enter an arm type. In this example, the first visit is considered naïve, whereas subsequent visits are informed. This distinction links the ALBs observed in the EPM test to foraging behaviours in wild-type rodents. Foraging behaviour encompasses any action taken to locate and exploit a food source. (Davidson et al., 2019; Owen-Smith et al., 2010). Foraging behaviour, particularly in new environments, evokes the conflict between safety and exploration that typifies rodent ALB in the EPM (Lima & Dill, 1990; McArthur et al., 2014). Theoretically, foraging decisions are modelled using Optimal Foraging Theory, which explains movement as the result of an attempt to maximise resource accumulation while minimising resource expenditure (Pyke, 2019; Stephens, 2008)

A key component of Optimal Foraging Theory is the Marginal Value Theorem, which predicts movement between resource patches (Charnov, 1976; King & Marshall, 2022; Pyke & Starr, 2020). Briefly, this theory predicts how long an animal should remain in a patch with resources when faced with deciding to stay or move to novel patches that may provide greater resources. For example, when that resource is food, optimal foraging occurs when minimal net energy is expended while maximal net energy is gained, which can only be achieved when the animal correctly predicts the resource value of the current patch and any possible patches to which it may travel. This decision obviously includes a degree of uncertainty that is proportional to the novelty of the possible destinations. Critically, modern theories of optimal foraging balance the risk of predation against the value of energy gain when assessing movement patterns (Found, 2022; Hintz & Lonzarich, 2018; Křivan, 1996).

Rodents signal a greater predicted predation risk by shifting from cautious exploration to behaviours such as surveillance, avoidance, freezing and escape. Theoretically, ALBs occur in the absence of a predator but are typified by the motivation to minimise the likelihood of an encounter (Hoffman et al., 2022; Mobbs et al., 2018). Given this definition, ALBs are considered a cost to net energy gain, resulting in suboptimal foraging. Thus, if the net energy of a foraging activity is a function of energy expenditure and energy gained, both are modified due to slower accumulation and increased non-foraging behaviours such as surveillance and caution.

The utility of NEG as a measure of ALB relies on the assumption that a rodent within the EPM is acting as an optimal forager in a novel environment. When a rodent anticipates a high threat risk, it will avoid exploring the area within open arms, as this region poses the highest risk of encountering a predator. When the expected risk of predation is low, rodents explore the available environment to evaluate possible foraging opportunities. Critically, the expected energy gain from the open arms is nonzero when the arms are novel to a naïve rodent, but this value diminishes commensurate with the actual lack of opportunities to forage once rodents are informed of this through exploration. Consequently, the relative expected net energy gain of open arm areas is lower than that of closed arms following exploration. This deficit exists because, while both arm types lack foraging opportunities, the closed arms provide greater proportional safety than the open arms, so fewer ALBs must be performed, reducing energy expenditure. This transition from naïve to informed and its impact on the value of the arm types means that an emphasis should be placed on the first visit to a novel location.

The transition from naïve to informed creates a dichotomy within the time spent in the open arms. Conventionally, time spent in the open arms considers all time spent within the open arms to have the same value across the testing period and to be generated by a single motivation. In contrast, we propose that the value of time spent in the open arms is nonlinear across time and is generated by two motivations: novel exploration and informed preferencing (see Fig 1A). This distinction between exploration and preferencing extends to when the rodent is not spending time in the open arms, creating four unique states throughout testing: avoidance, exploration, not preferencing, and preferring (Fig 1B). These first two states capture the motivational conflict that governs the assessment of anxiety-like behaviour during exposure to the EPM. Consequently, they form the basis for what is measured using NEG (Fig 1C).

**Figure 1.**
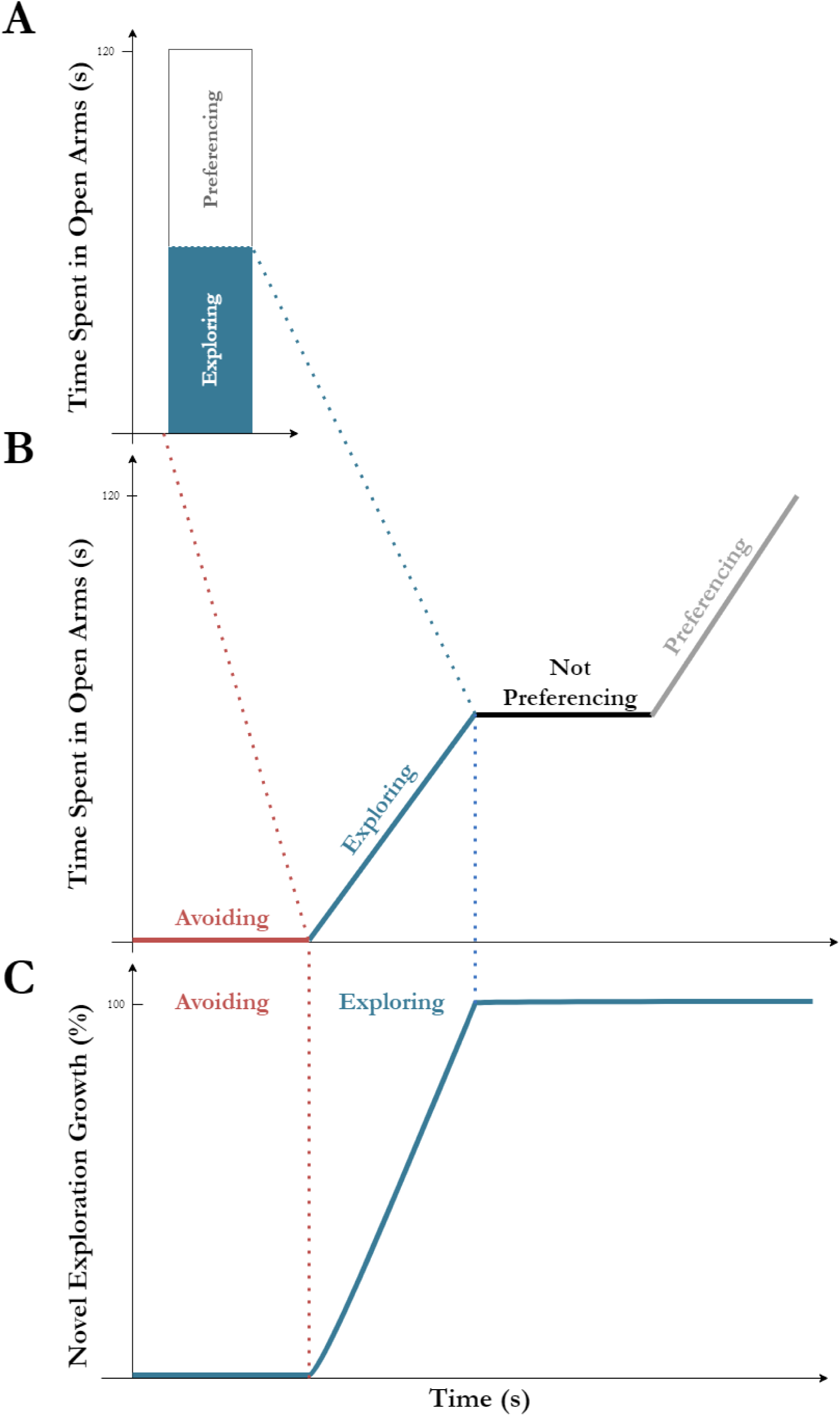
Mapping of time spent in the open arms of the EPM to novel exploration growth. The time spent in the open arms comprises two qualitatively different states, exploration and preferencing (A), where exploration represents the first visit to an area and preferencing represents subsequent visits. These two states form part of a larger suite of behaviours when measuring time spent in the open arms throughout testing. This suite includes avoidance of the open arms, which is distinct from not preferencing them once they have been explored (B). Given their relevance to anxiety, avoidance and exploration of the open arms are mapped onto a growth curve of novel exploration via NEG (C). Avoidance is inferred from a lack of initial growth in the curve, whereas exploration is evident from the phase of growth.

NEG growth curves will differ between individual rodents as they vary in their level of anxiety-like behaviour. Nonetheless, these curves tend to match one of three distinct behavioural phenotypes: Exploratory, Delayed and Avoidant. As shown in Figure 2, a rodent exhibiting an Exploratory behavioural phenotype traverses the open arms during the first phase of exploration, either exclusively or mixed with exploration of the closed arms. This pattern contrasts with an Avoidant phenotype, where the rodent exclusively explores the closed arms, indicating a preference for safety. Critically, a rodent exhibiting the Delayed phenotype treats the arms differently over time, displaying an initial avoidance of the open arms that transitions into an exploration of them. Although subsequent exploration of the open arms could occur in multiple phases, the key indicator of this phenotype is whether this exploration reliably occurs after a period of preferencing the closed arms. This preferencing causes a plateau in NEG, showing that exploration has ceased and that safer areas are preferred, indicating a higher overall ALB.

**Figure 2.**
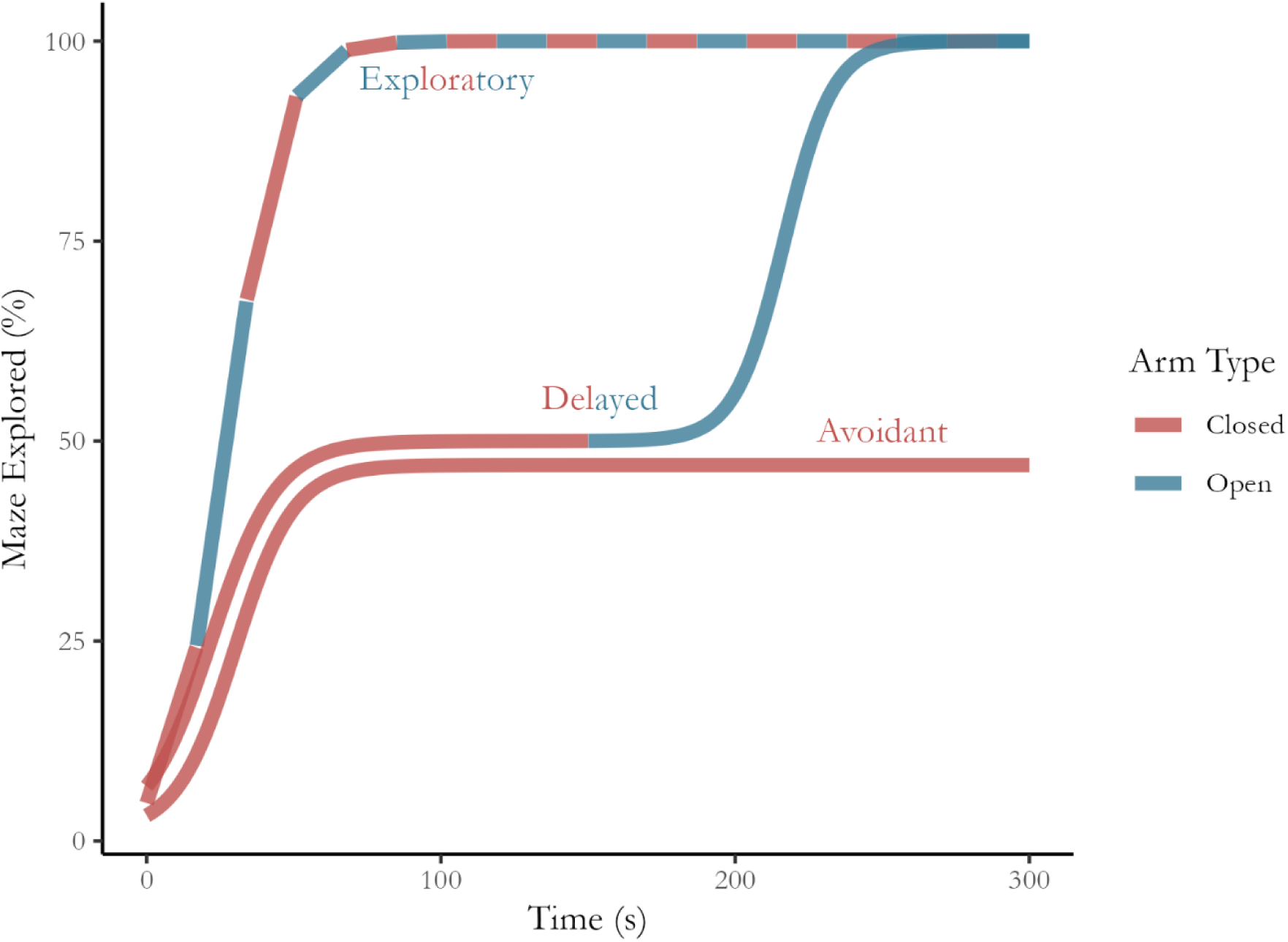
Exploratory, Delayed and Avoidant exploration phenotypes. The three behavioural phenotypes can be separated by their differences over time and by the extent of open arm exploration. The line colour indicates which arm type is being explored during testing. Note: For illustrative purposes, the lines have been separated during the first 20 s across all phenotypes and up to 200 s for the Delayed and Avoidant phenotypes.

The growth curves in Figure 2 highlight that to infer the level of ALB, there is a need to estimate both the phases of exploration and the proportion of the total area of the open arms that were explored. As outlined above, assessing the phases of exploration establishes whether a preference existed between arm types. If the closed arms were preferred earlier than the open arms, this indicated a high level of ALB. In addition, the amount (i.e., area covered) of open arm exploration is important because a low amount indicates a lack of exploration depth. This lack of commitment to physically exploring the open arms of the EPM indicates a high level of ALB, particularly if this amount of exploration is much lower than the exploration of closed arms.

Using the heuristics outlined above, Figure 3 shows how the exploratory behavioural phenotype is mapped to a gradient of ALBs by assessing the phases of exploration and the difference between the amount of exploration observed in each arm type. An Avoidant phenotype is inferred when no open arm exploration is observed and only the closed arms are explored, indicating the highest level of ALB. A Delayed phenotype displays a similar initial phase of closed arm exploration, but subsequent phases include open arm exploration. This sequence can be interpreted as indicating moderately-low to moderately-high ALB, depending on the extent to which the open arms were explored compared to the closed arms. Conversely, an Exploratory phenotype must explore a mix of arm types or the open arms exclusively within the first exploration phase. This caveat exists because the critical element distinguishing this phenotype is the lack of initial avoidance of the open arms. The absence of avoidance indicates that exploration, not safety, motivates this phenotype’s traversal in this phase. Critically, the level of ALB inferred from this sequence still depends on the extent to which the open arms were explored compared to the closed arms. Low ALB is inferred where the amounts are similarly high; however, if the open arms are explored at a much lower comparative rate, then higher ALB is observed.

**Figure 3.**
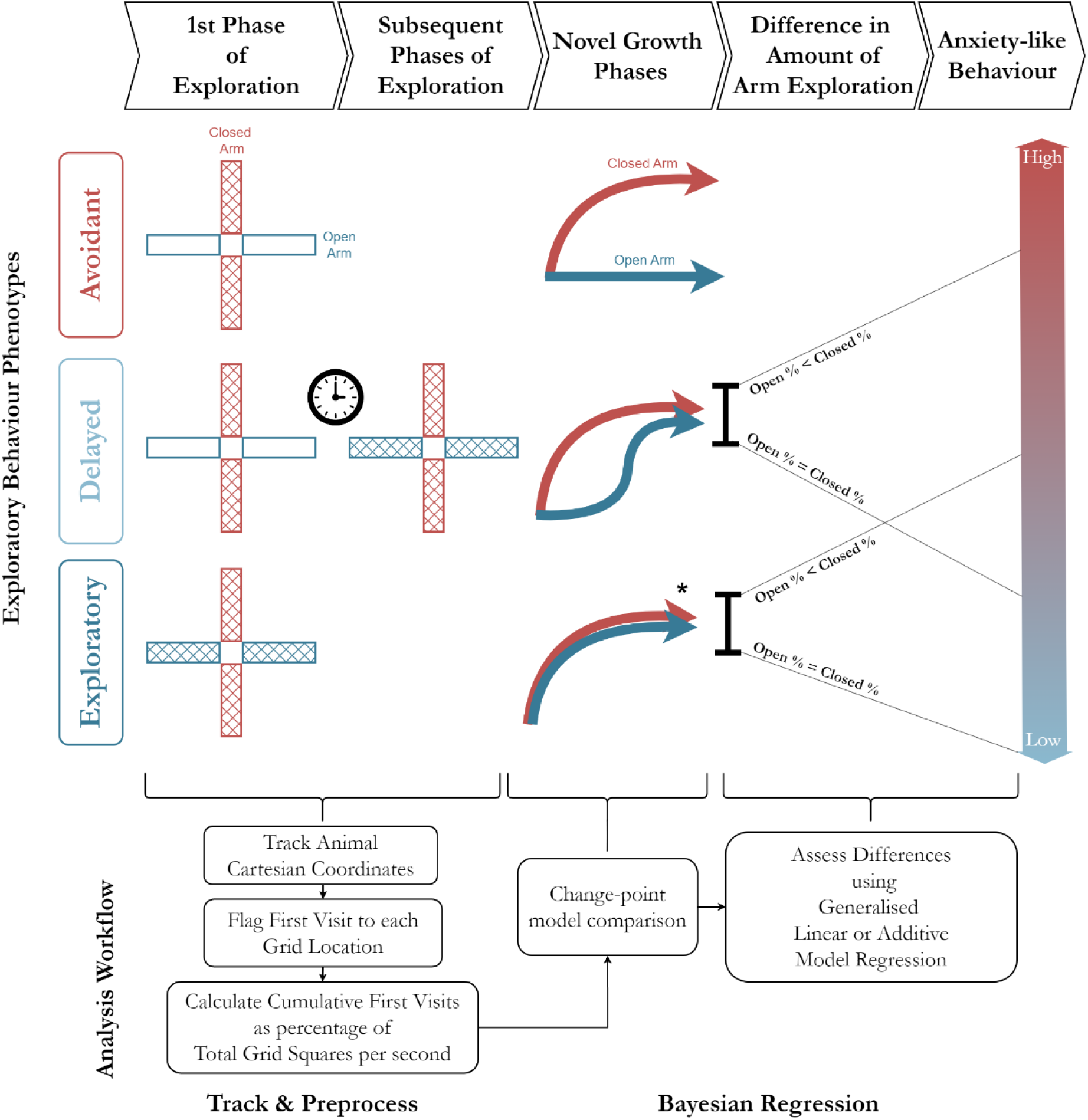
Mapping the exploratory behavioural phenotype to the ALB via novel exploration growth, including the analysis workflow. The position of a rodent in the maze is tracked and preprocessed to enable a temporal analysis of exploration (left to right from the 1_st_ phase to subsequent phases). The cumulative first visits for the total maze and by arm type are evaluated via change-point models to establish their “Novel Growth Phases”. The total amount of exploration was then compared between arm types to establish how much ALB was observed.

Both conventional measures and NEG emphasise physical commitment to the open arms; however, NEG extends this emphasis by examining the depth and timing of commitment. When a rodent delays or abstains from exploring the open arms, this is considered to indicate that the safety of the closed arms provides more perceived benefit than the novelty of the unexplored open arms. Critically, a rodent demonstrates an exploratory phenotype whether its first phase of exploration mixes arm types or whether it explores only the open arms. This is because an exploratory rodent would treat the arms equally as novel areas and explore them agnostically. Conversely, a rodent displaying a greater ALB would show a preference for the closed arms, which would manifest as multiple exploratory phases where exploration of the open arms was delayed compared to that of the closed arms.

Casarrubea and colleagues (2013) claim that sequencing behaviours in time during testing in the EPM is critical to its construct validity. However, recent models ignore how behaviours are interrelated over time. (Arantes et al., 2013a; Tejada et al., 2010). This oversight makes them incompatible with the dramatic changes that occur after information has been gathered and marginal value drops. This issue is further magnified by the lack of exploitative opportunities in the EPM, such as the presence of food, rendering the value of the open arms largely limited to their initial novelty. Conversely, the inferential power of NEG in estimating the level of ALB is made possible through a simple decision framework using linear and nonlinear time series models.

### Method

The number of exploratory phases and NEG were estimated and compared using a multistage workflow (see “Analysis Workflow” in Figure 3). The workflow comprises two discrete sections: data collection and preprocessing to calculate NEG and Bayesian regression analysis to assess the change points in the time series and conditional effects of interest, such as the effect of phenotype, arm and phenotype-by-arm type interaction effects.

### Data Collection and Preprocessing

First, the position of the rodent’s centre of mass is tracked per second and translated into cartesian coordinates, with the centre of the maze being assigned as the origin (0,0). Various standalone programs, such as EthoVision and ANY-maze, can be used to track movement, as can packages such as pathviewr and TrackPy in R and Python. The remaining steps in the workflow were conducted in R using custom-developed scripts.

The coordinates of the rodent are then mapped to a 1x1 cm reference grid using Cartesian coordinates with the origin at the centre of the maze. Each square in the reference grid is allocated a unique identifier, which enables each visit to a square by the rodent to be recorded in time (see supplementary figure S-1). This process requires single-point tracking of the animal. Although more accurate than multipoint tracking, this method can neglect the exploration attributable to the head, which is often used near edges such as those in the open arms (Becker et al., 2023; Chetverikov & Verestoy, 1998). Critically, to accurately track the animal, the sampling rate of the tracking video must be high enough to map movement into each square. A rate of 25 frames per second is recommended.

For 300 seconds, the cumulative percentage of the total maze or specific arm type explored is computed by aggregating the flagged grid squares per second. When expressed as a time series, this cumulative measure depicts the temporal phases in which this traversal occurs and the depth to which areas are explored during these phases via the percentage value.

### Comparing Phase Models using Change-Point Regression

Bayesian change-point regression analysis is conducted on NEG over time for each phenotype’s total maze exploration and by arm type to evaluate exploratory phase dynamics. Change-point models are fitted using the mcp package in R, which produces inferences regarding the location of changes in the trend of a time series (Lindeløv, 2020). Like other bounded exponential growth phenomena, this time series is modelled using an *n*-sized sigmoid function (Kucharavy & De Guio, 2015). As shown in Figure 5, single and dual growth phase sigmoid functions are used to model change points in the NEG time series (see Supplementary S-2 for model specifications). In practice, the NEG time series for the entire maze and by arm type are modelled individually. The use of individual time series allows for an overall inference to be made regarding exploration and arm type-specific conclusions.

**Figure 5.**
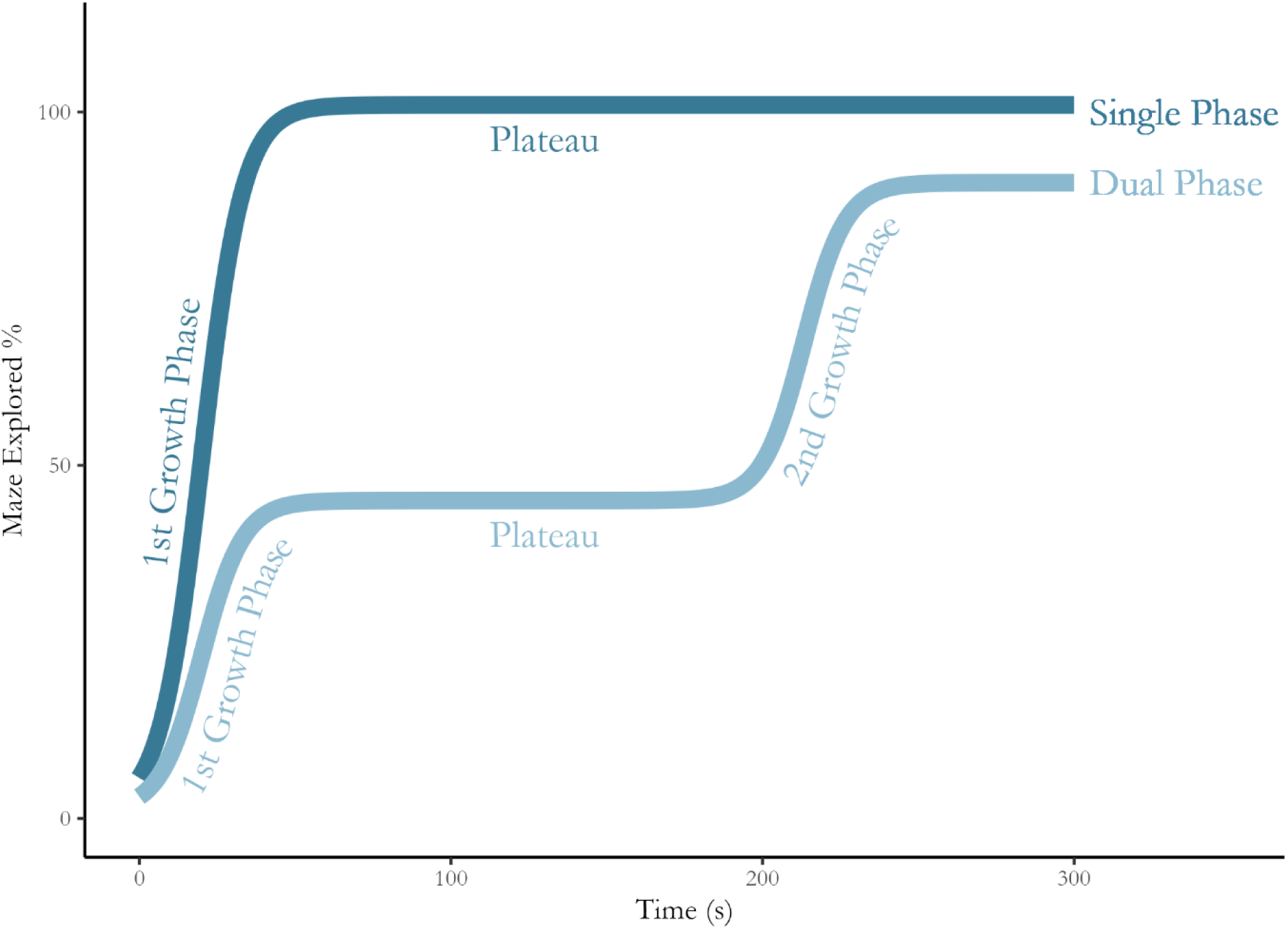
Single- and dual-phase exploration models for the maze. Solid coloured lines indicate novel exploration growth in the EPM over time for single- and dual-phase models. Both models show initial growth that plateaus with the dual-phase model characterised by a second growth phase.

### Estimating Marginal and Conditional Effects using Bayesian Regression

The current methodology follows the recommendations made by Betancourt (2020) when conducting Bayesian analysis for both change-point and generalised linear regression components of this workflow. Prior predictive checks are performed to assess the appropriateness of the chosen prior distributions. These checks involve generating data from the prior predictive distribution and evaluating whether these data are consistent with known or plausible values in the context of the study. Posterior predictive checks are performed to evaluate the model’s fit and its ability to reproduce observed data patterns. These checks involve visually comparing the observed data with data simulated from the posterior predictive distribution to assess model adequacy.

Model fit is also verified by ensuring that the parameter *R̂* (Rhat) values are within the tolerance range of [0.9,1.1] and that the effective sample size (ESS) is greater than 1000 (Bürkner, 2017). Finally, leave-one-out cross-validation using Pareto-smoothed importance sampling (LOO_PS_-CV) is conducted using the loo R package to ensure adequate model fit (Vehtari et al., 2022). LOO_PS_-CV evaluates model fit by estimating the expected log posterior predictive distribution (elpd_loo_) and the effective number of parameters used to fit an out-of-sample dataset (Vehtari et al., 2017).

Change-point model comparisons are also made via LOO_PS_-CV. If both single- and dual-phase models meet the predictive checks and convergence criteria, models are ranked according to the highest elpd_loo_ with the lowest number of parameters. The model that returns the higher elpd_loo_ is preferred; when elpd_loo_ is equivalent, the most parsimonious model is selected.

Once phase model selection has been completed, the effects of interest are analysed using Bayesian mixed model regression. For the simulation below, the effects of interest include the marginal phenotype effect for the total maze and the conditional effect of arm type. Generalised linear mixed models can be used to estimate the effects of interest when growth curves of interest are best captured as single-phase growth curves. If any time series is best captured by a multiphase growth curve, such as the dual phase model specified above, then Bayesian generalised additive model regression is applied to estimate effect characteristics. These models are also fit via the brms package using smooth functions from the mgcv r package (Wood, 2011).

Generalised additive models are chosen because, as a semiparametric specification, they can model nonlinear relationships between the response and predictor variables very flexibly while still producing interpretable effect estimates via smoothing functions (Militão et al., 2023; White et al., 2020). The smoothing functions can be fitted componentwise, for example, per phenotype or phenotype/arm type interaction, and are specified using a smoothing spline. For this workflow, a thin plate smoothing spline is chosen because it has excellent multidimensional properties, making it useful for assessing spatiotemporal foraging patterns (Christensen-Dalsgaard et al., 2017; Preisler et al., 2004).

Bayesian mixed model regression analysis is conducted using the brms r package (Bürkner, 2018), with marginal and conditional effects being estimated using the BayestestR and emmeans packages or using custom functions (Lenth, 2022; Makowski, Ben-Shachar, & Lüdecke, 2019). Marginal and conditional effects are evaluated inferentially using the SeXit framework, which reports estimates for centrality, uncertainty, existence, and significance (Kruschke, 2018; Makowski, Ben-Shachar, Chen, et al., 2019; Schwaferts & Augustin, 2020).

Centrality and uncertainty are reported using the median and highest density interval (HDI) of the distribution of the effect. The effect’s existence is indicated by assessing the probability of its direction (PD). In contrast, significance is established by constructing a region of practical equivalence (ROPE), which verifies when an effect is negligible because it is practically equivalent to zero. To make the language synonymous with null hypothesis significance testing, the effect is reported as confirmed and equivalent to zero when greater than 97.5% of the posterior remains within the ROPE. Conversely, the equivalence of the effect to zero is rejected when less than 2.5% of the posterior remains within the ROPE. In cases where more than 2.5% but less than 97.5% of the posterior distribution of the effect remains within the ROPE region, the practical equivalence to zero is reported as undecided. For further details on constructing a region of practical equivalence, see Kruschke (2018) and Schwaferts and Augustin (2020).

### Simulating Phenotypical Exploratory and Conventional Behaviours

A simulation study was conducted to demonstrate how the change-point model comparison and regression analysis elements of the NEG workflow can be applied to samples of interest. Specifically, this study analyses the total maze and arm-specific NEG for the three behavioural phenotypes proposed above to show how to use NEG to infer the level of ALB being observed. The exploratory behaviour for the entire maze was simulated using the double sigmoid function with a sigmoid per arm type (see Supplementary Figure S-3 for parameter values for growth rates and inflection points by phenotype for the total maze by arm type). Parameter values were chosen because they produce reasonable exemplars for the expected time series for each phenotype. The Exploratory phenotype is expected to show rapid, arm-agnostic exploration growth at test onset; thus, arm growth rates are sampled from the same distribution, as are the inflexion time points for each arm. Conversely, the Delayed phenotype is expected to show an earlier inflection time point for the closed arm than for the open arm. Finally, the growth rate and the inflection points will differ between arms for the Avoidant phenotype.

Figure 6 visually depicts the simulated NEG by phenotype for the total maze (A) and by arm type for all three phenotypes (B:D). As expected, the Exploratory phenotype showed greater total exploration growth due to being arm-agnostic (B), whereas only closed arms were initially explored by the Delayed (C) and Avoidant phenotypes (D). After 200 s of testing, the Delayed phenotype fully explored the open arms, reaching an equivalent total NEG to the Exploratory phenotype. These NEG time series also included three conventional ALB measurements, latency to enter the open arms, number of open arm entries and total time spent in the open arms, according to the following criteria:

- The average velocity is assumed to be identical regardless of the arm type.
- The first entry into the open arms is followed by full exploration of that arm type.

**Figure 6.**
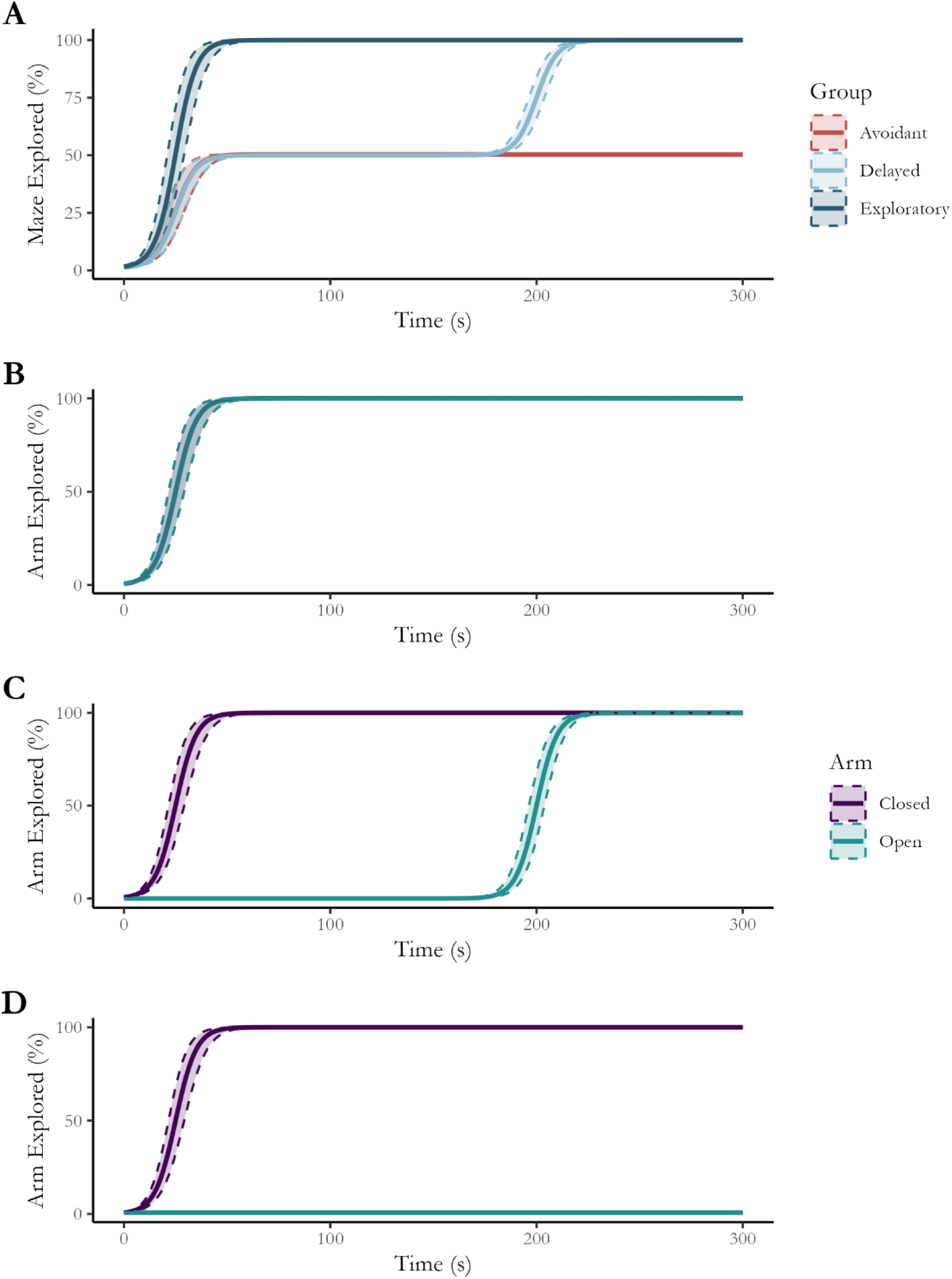
Total maze exploration and exploration by arm type per phenotype. Exploratory and Delayed phenotypes initially showed similar rates of exploration growth for the entire maze (A); however, while the Exploratory phenotype (B) showed an overlap between NEGs by arm type, Delayed (C) and Avoidant (D) phenotypes displayed differences over time due to their subsequent exploration or complete avoidance of the open arms.

These assumptions ensure that the conventional ALB behaviours faithfully represented the growth curves and did not create artificial differences between phenotypes and arm types. Basal locomotor activity is typically operationalised by measuring average velocity, where outliers may be removed from analysis due to illness or malaise (Schrader et al., 2018). Therefore, keeping this constant between exploratory phenotypes removes any bias associated with locomotive patterns. In addition, the requirement that an arm type is fully explored following the first entry ensures that no bias is introduced when estimating the number of entries or time spent within an arm type. Although we expect real data to include much more variance in the NEG time series they generate, these phenotypes demonstrate an idealised version of exploratory, delayed and avoidant behaviours. Subsequent publications will show that real-world observations can be reliably characterised using one of these phenotypical behavioural patterns and analysed using the proposed workflow.

Given these criteria, the minimum number of open arm entries required by the Exploratory and Delayed phenotypes is two, with time spent in the open arms also being equivalent. Conversely, the Exploratory phenotype enters the open arms early in testing, whereas the Delayed phenotype enters midway through testing. In contrast, the Avoidant phenotype does not enter the open arms at all, so the latency to enter is set to the maximum testing duration, with zero entries into the open arms being observed and no time spent within them.

The distributions by phenotype for each open arm measurement value were simulated 1000 times using a truncated Gaussian distribution between 0 and 300 since, in the case of the time spent in the open arms and the latency to enter them, the values must be positive real numbers that do not exceed 300 (see supplementary figure S-4 for simulation value summary and generation distribution).

## Results

### Conventional Open Arm Measures

Bayesian linear regression was conducted using phenotype as a predictor to evaluate the effect of phenotype on conventional open arm measures. Default priors were used for all parameters, and all models were well specified. Figure 7 shows each conventional measure’s mean and variance by phenotype, including the equivalence decision. As expected, the latency to enter the open arms of rodents with an Avoidant phenotype, total entries and time spent within the open arms were not practically equivalent to the values measured using these parameters for Exploratory or Delayed phenotype rats. Conversely, only the latency to enter the open arms was not practically equivalent between the Delayed and Exploratory phenotypes (see supplementary figure S-5 for effect summary and SeXit reporting table).

**Figure 7.**
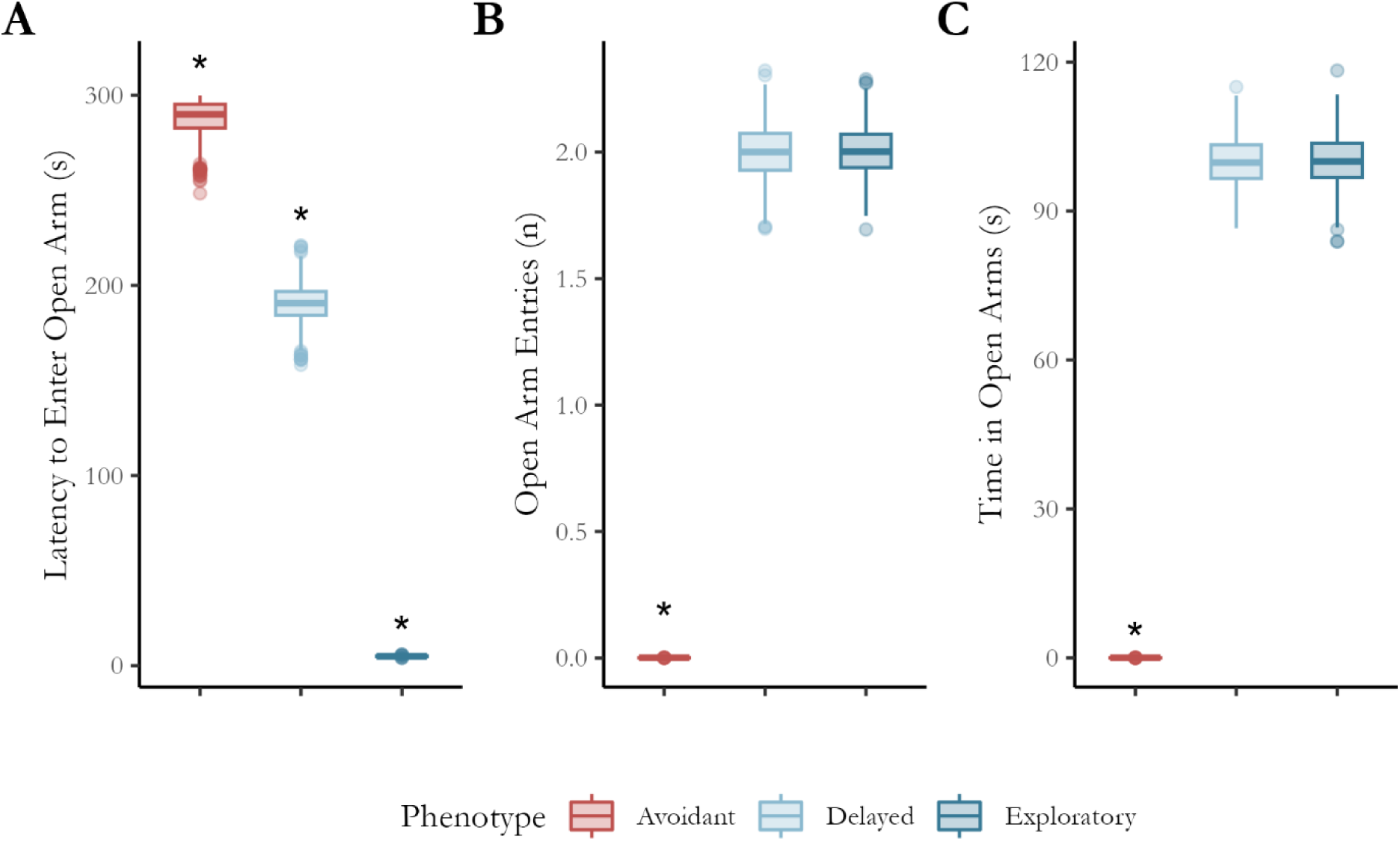
Mean and variance statistics for open arm preferencing metrics. The three phenotypes demonstrated differences in their latency to enter the open arms (A); however, only the Avoidant phenotype displayed differences in the number of entries (B) into and time spent within (C) the open arms. An asterisk (*) indicates that it is not practically equivalent to any other phenotype.

### Estimating Novel Exploration Phases via Change Point Regression

To determine whether NEG occurred in a single or dual phase, two change-point regression models were fitted per phenotype to illustrate movement across the entire maze (see Supplementary Figure S-2 for full model specifications, including priors, and S-6:7 for prior predictive checks).

The posterior distributions and model fit diagnostic criteria for all phenotypes can be found in the supplementary materials (see S-8 in the supplementary materials). Using the diagnostic criteria outlined earlier, we determined that the single-phase model is the preferred choice for total maze NEG for the Exploratory and Avoidant phenotypes, whereas the Delayed phenotype is best modelled using a dual-phase growth model.

Comparing NEG between Phenotypes for the Total EPM Given the heterogeneity of preferred phase models across phenotypes, a generalised additive mixed effects regression model was used to assess the effect of phenotype on NEG for the total maze (see Figure S-9 in the supplementary materials for full model specifications, including priors and S-10 for posterior predictive checks and model diagnostics).

Figure 8 shows the posterior of the marginal effect of phenotype over time for the total maze. The NEG for the Exploratory phenotype is greater than that for the Avoidant (A) and Delayed (B) phenotypes, although this latter difference diminishes after 200 s. Conversely, the exploration growth of the Delayed and Avoidant phenotypes (C) is equivalent for the first 200 s of testing but diverges due to the phase of open arm exploration by the Delayed phenotype (see Figure S-11 in the supplementary materials for a full regression summary table).

**Figure 8.**
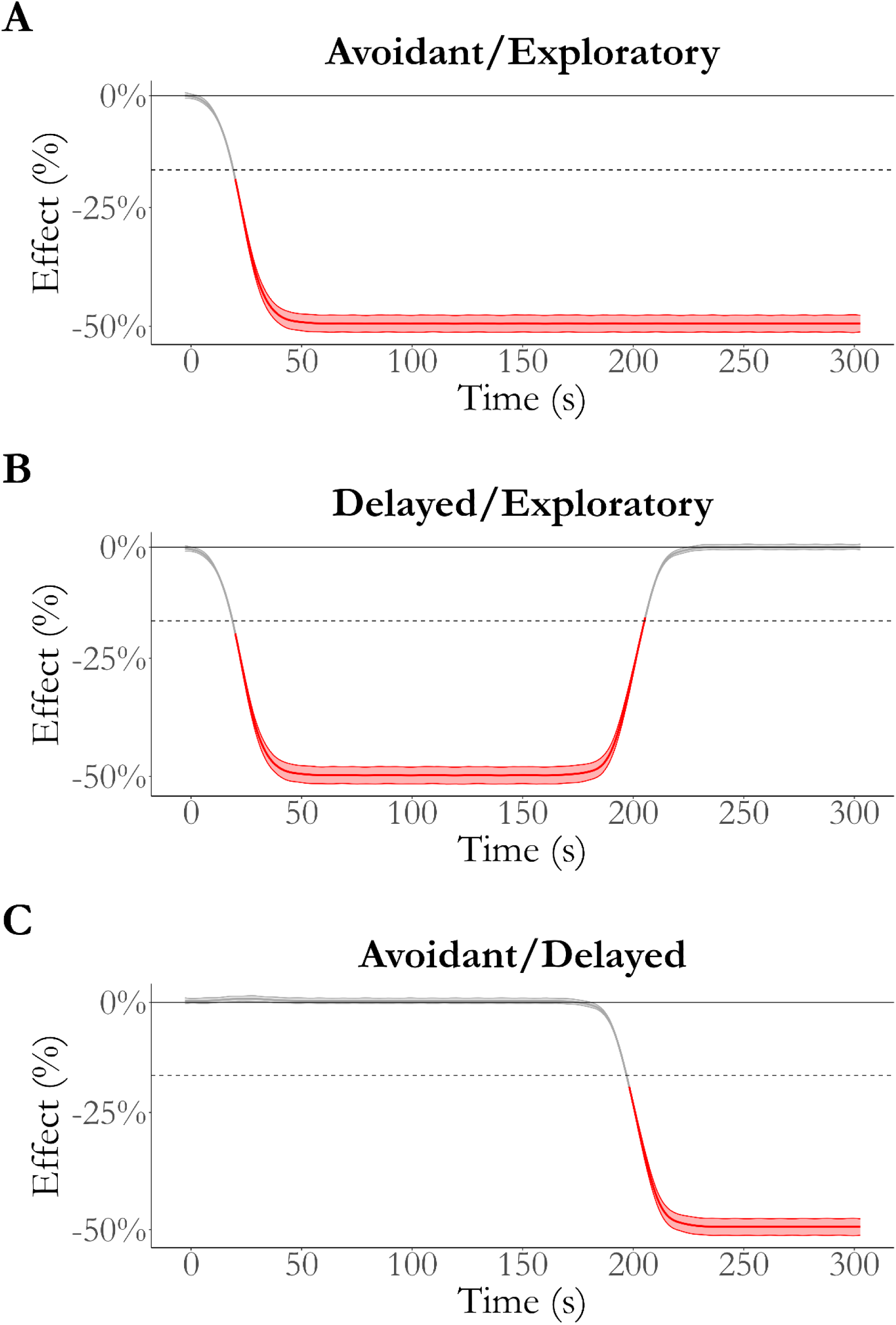
Effect of phenotype over time on total maze NEG. The Avoidant and Exploratory phenotypes (A) are not equivalent to zero beyond the first few moments of the testing, whereas the difference between the Exploratory and Delayed phenotypes (B) is not equivalent for the first 200 s. The difference between the avoidant and delayed phenotypes is equivalent to zero for the first 200 s of testing (C). Red indicates that the effect is not practically equivalent to zero. Note: ROPE Region = 0 ± 18%, as indicated by the dashed line.

### Comparing Arm Types between and within Phenotypes

To determine whether NEG by arm type for each phenotype occurred in single or dual phases, the change-point regression used for the total maze exploration was applied to these contrast pairs (see s-12 in the supplementary materials for posterior predictive checks and model diagnostics). As expected, all phenotype/arm exploration contrast pairs were best modelled as single-phase growth curves.

Given that all contrast pairs were best modelled using a single phase of exploration growth, a generalised linear mixed model was used with a binomial link function to assess conditional differences in hierarchical data such as subject-level time series (see Figure S-13 in the supplementary materials for a full model expression including priors and S-14 for a regression summary table including diagnostic values).

Figure 9 shows the effect of arm type over time between each phenotype. As expected, the conditional effect of arm type rapidly became negative for the Avoidant phenotype (A). Conversely, the effect of arm type is always equivalent to zero for the Exploratory phenotype (B) and becomes equivalent to zero for the Delayed phenotype in the later stages of testing (C). Figure 10 shows the effect of phenotype on open arm NEG, where the difference between the Avoidant and Exploratory phenotypes rapidly reached 100% (A). Conversely, the difference between Delayed and Avoidant (B) became practically significant only after 185 s of testing. In contrast, the Delayed and Exploratory phenotypes are practically equivalent only after 200 s of testing after being significantly different for the majority of testing.

**Figure 9.**
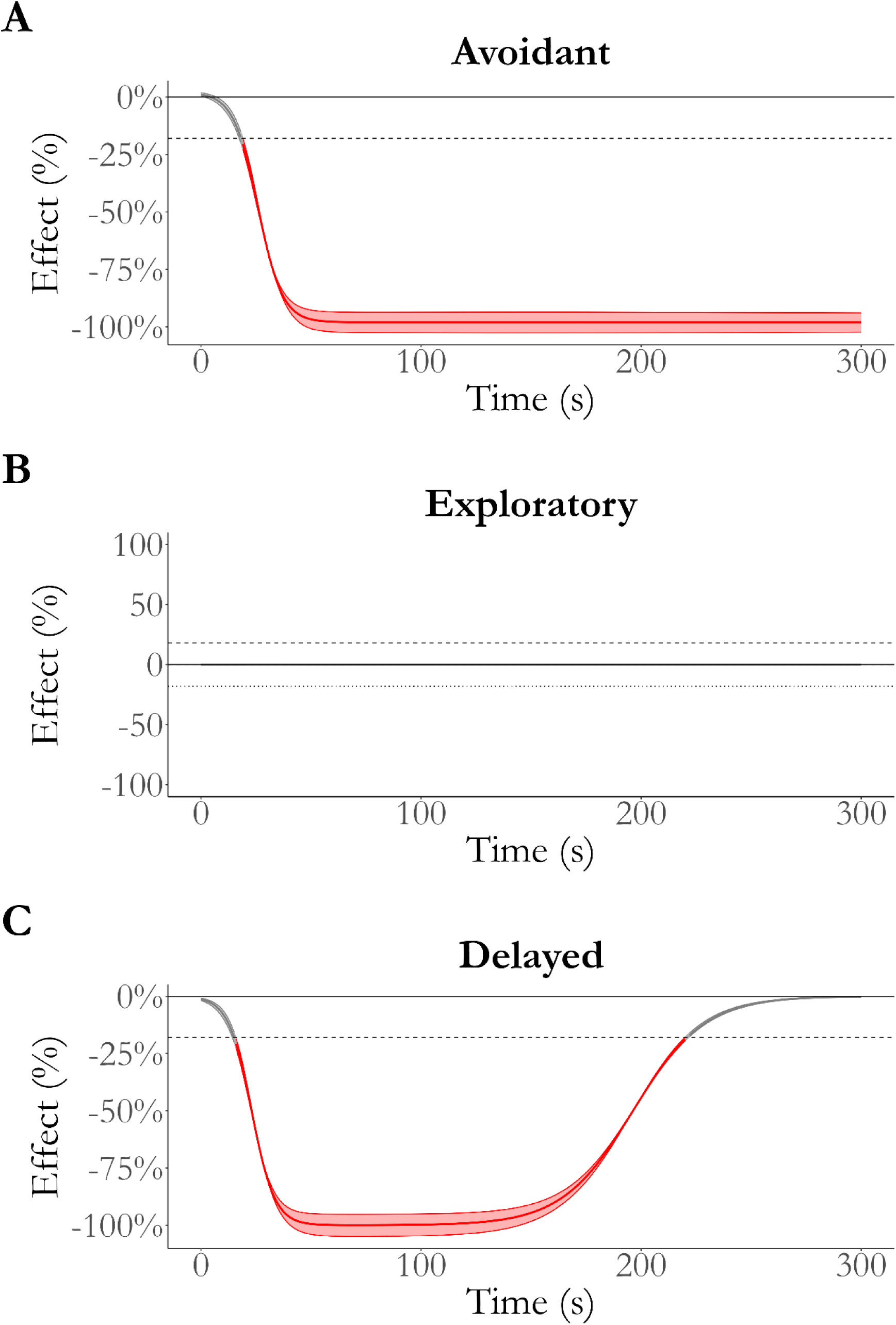
Effect of arm per phenotype over time. The difference between closed and open arm NEG for the Avoidant phenotype (A) quickly reaches 100%, which contrasts the effect for the Exploratory phenotype (B), which is practically equivalent to zero for the duration of testing. Finally, the effect of arm type for the Delayed phenotype initially reaches 100% but reduces back to practical equivalence after ∼ 225 seconds. Red indicates that the effect is not equivalent to zero, where the thick line indicates the median effect over time and the thinner lines indicate the region of the full posterior of the effect. ROPE Region = 0 ± 18%, as indicated by the dashed line.

**Figure 10.**
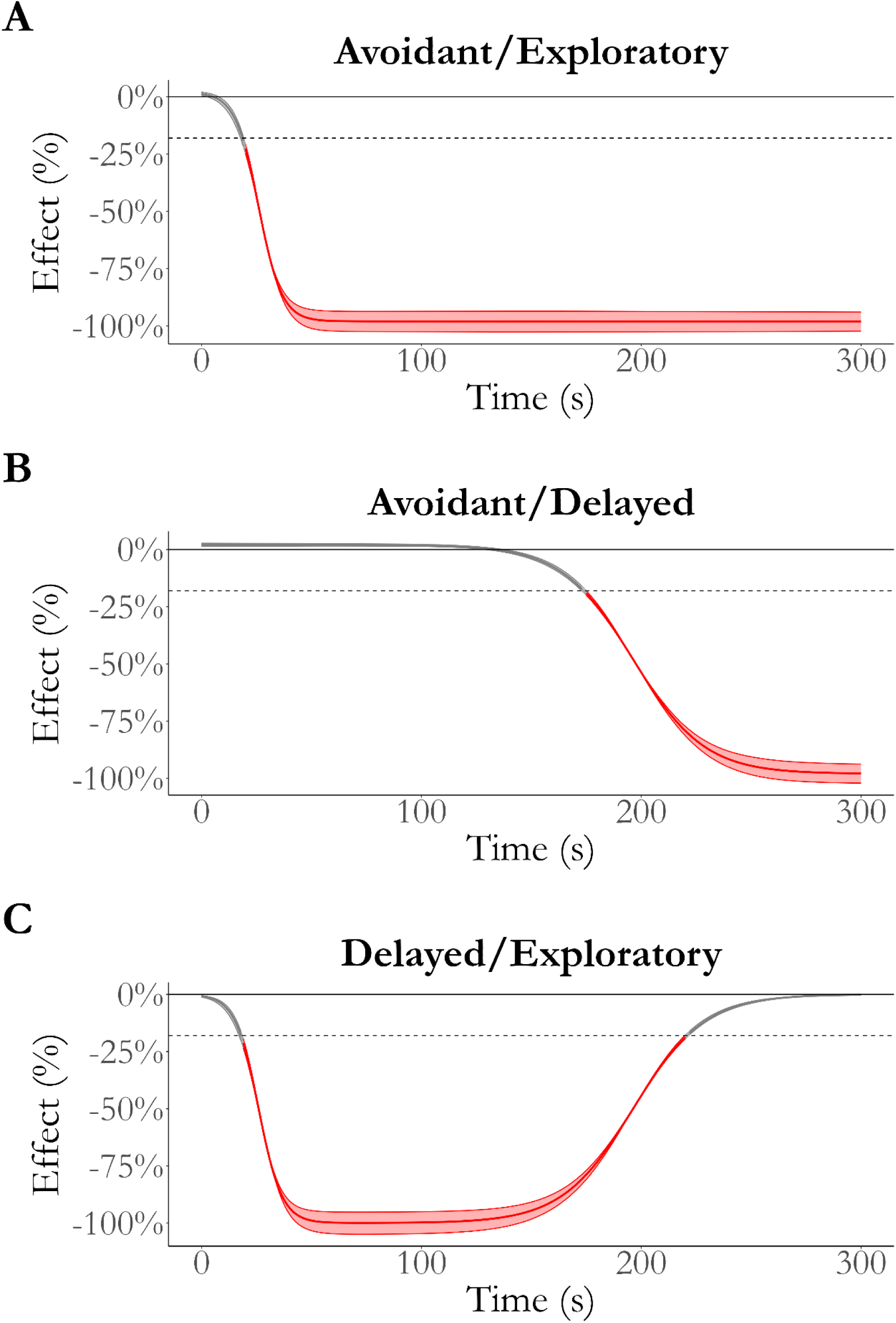
Effect of phenotype on Open Arm NEG over time. The difference between the Avoidant and Exploratory phenotypes (A) quickly reaches 100%, while the effect reaches maximum later when compared to the Delayed phenotype (B). Finally, the difference between the Delayed and Exploratory phenotype initially reaches 100% but reduces back to practical equivalence after ∼ 225 seconds. Red indicates that the effect is not equivalent to zero, where the thick line indicates the median effect over time and the thinner lines indicate the region of the full posterior of the effect. ROPE Region = 0 ± 18%, as indicated by the dashed line.

## Discussion

This paper aimed to provide a single, coherent measure of exploratory and avoidant behaviours of rodents in the EPM. To achieve this, we propose evaluating novel exploration growth over time across the entire area of the maze and within each arm type. By simulating three distinct exploratory behavioural phenotypes, we showed that NEG uniquely and coherently maps movement throughout the maze to levels of ALB. These inferences were produced by analysing the phases of traversal of rats using change-point analysis and comparing these phases using generalised additive regression.

According to the NEG analysis, the simulation of the Exploratory phenotype captured the least anxious behavioural characteristics since the change-point analysis indicated that exploration took place in a single arm-agnostic fashion. The simulation of the Delayed phenotype appeared marginally more anxious behaviourally compared to the Exploratory phenotype, as it displayed two, arm-specific phases of exploration that signalled initial avoidance and caution that receded over time. This reduction in exploratory activity reflects a dynamic updating of threat imminence over time via information gathering that lowers avoidance and promotes exploration of the open arms (Davidson et al., 2019; Nonacs, 2001). The simulation of the Avoidant phenotype displayed maximal ALB since it displays a continuous aversion to the open arm regions of the EPM. This continual aversion indicates that the naïve predicted value of the open arms never exceeds that predicted value of the safety provided by the closed arms.

Theoretically, a higher ALB is inferred when familiar, safer areas are preferred, while novel, riskier areas are avoided (Montgomery, 1951; Walf & Frye, 2007). In the EPM test, this behavioural pattern is partially observed by measuring the latency to enter the open arms, as it directly captures the avoidance of the riskier areas; however, it cannot substantiate the preferencing of safer ones. Despite this, the latency to enter the open arms was still able to correctly indicate that the Delayed phenotype displayed higher ALB than the Exploratory phenotype. Conversely, the NEG purposefully identifies when closed arms are not only being explored but are also preferenced prior to an initial entrance into the open arms. An example of preference is depicted by the Delayed phenotype’s effect on NEG between arm types, where the time series showed that exploration growth plateaued before exploration of the open arms.

In contrast to the latency to enter the open arms, the conventional measure of the total number of entries into the open arms could not differentiate between the levels of ALB displayed by the Delayed and Exploratory phenotypes using their respective simulated values. Instead, analysis of this measure erroneously indicated that the two phenotypes were practically equivalent. Notably, this failure would become misleading if the Delayed phenotype made even one additional, shallow entry into the open arms prior to exploring them. In this hypothetical case, all three conventional measures would lead to orthogonally different interpretations: latency to enter the open arms indicating that the Delayed phenotype was more anxious, while the total time spent in them indicating no difference in anxiety-like behaviour and the total number of entries indicating lower anxiety-like behaviour. Critically, there is no justifiable way to select the optimal parameter from the three conventional measures because they evaluate different facets of ALB (Cruz et al., 1994; Walf & Frye, 2007).

The current study demonstrated that measuring physical exploration across time can isolate early avoidance from eventual exploration. This strength was evidenced by successful identification of the dual phases of the simulated NEG across the entire maze by the Delayed phenotype. Critically, when evaluated using NEG trends, an early, shallow entry into the open arms is correctly distinguished from the true, deep exploration displayed by the Exploratory phenotype. In doing so, NEG, as a single measure, can remedy inconsistencies imposed by using the three conventional measures.

A principal limitation of the current methodology is that the speed of movement impacts the calculation of NEG when above a certain rate. When a rat is moving very quickly, typically at speeds greater than 25 cm/s, resolving the areas it has entered becomes difficult. At these speeds, novel exploration estimation requires higher resolution grids and frame rates for accurate tracking. Critically, this lack of resolution of the animal’s position is justified since the faster the animal moves, the less information it can gather about any individual location. Given the ethological basis for exploration is as an information gathering behaviour, the speed of traversal through an area should vary proportionally with the amount of information gathered about the locale. Therefore, while NEG estimation does carry a bias toward slower exploration, this reflects greater certainty that a location has been explored compared to higher movement speeds. This assertion has been validated during testing, where speeds above 25 cm/s captured at 24 frames per second do not impact overall trends in real-world datasets. Despite this, end users should be aware that such speeds may produce small variances in expected NEG values compared to the actual calculated values. This limitation also underscores the need to capture animals at an adequate frame rate to minimise the impact.

The current framework does carry limitations, however, particularly in the implementation of the regression analysis. For example, while generalised additive models are flexible enough to handle noisy real-world data, they require stronger assumptions about how these temporal patterns map to ALB. In particular, the choice of smooth function and number of basis functions impacts the fit and interpretation of the output and its generalisability (Ezhov et al., 2018; Harmening & Neuner, 2016). In contrast, the authors explored the use of fully specified sigmoid functions, which can make interpretation easier and more generalisable, but also leads to label switching when fitting (Ngah et al., 2016; Park et al., 2019). The effectiveness of this approach has been tested against real-world data, which will be presented in a series of follow-up publications, where NEG was able to isolate differences in ALB due to treatment. Additionally, the methodology will be further validated by users who apply it to data representing previously established anxiolytic interventions, such as benzodiazepines and antidepressants (Hogg, 1996; M. Kurt et al., 2000).

An essential limiting facet of NEG is that it only requires that users explicitly model the initial avoidance and exploration of an area or arm type. Preferencing can be inferred indirectly by comparing differences in growth between arm types, as shown by the plateau between exploration phases demonstrated by the Delayed phenotype. Further extensions could introduce more complex models that directly account for both exploratory and preferencing states, such as using partially observable Markov decision processes to include memory and agency in the model. Additionally, this proposed approach emphasises physical displacement and presence as critical indicators of anxiety during EPM testing. Future studies may incorporate dynamic Bayesian networks that include non-locomotor behaviours such as grooming or rearing to increase the resolution of the multidimensional presentation of anxiety during exposure to the EPM. Finally, the technical aspects of the workflow may pose a barrier to implementation for some users of the EPM. To combat this, the code used in this workflow is available, and frequentist alternatives to key analytical elements are easily substituted if the user prefers them.

## Conclusions

The present study demonstrated that novel exploration of the EPM accurately distinguished between three phenotypical anxiety-like behaviours. This capability was achieved using Bayesian change-point and generalised additive regression to model rodent behavioural data from EPM assays. The study also demonstrated that conventional ALB measures produce inconsistent and erroneous inferences when analysed using simulated data representing phenotypical exploratory behavioural patterns.

## Acknowledgements

We thank Dr. Antonina Govic for her assistance in reviewing the theoretical aspects of the proposed methodology. We would also like to thank RMIT University for providing ongoing non-financial support for the project. We would like to thank Epigenes Australia for funding the article processing charge for this manuscript to allow it to be published as open access.

## Declarations

### Ethics approval

Not Applicable

### Consent to participate

Not Applicable

### Consent for publication

Not Applicable

### Availability of data and materials

Not applicable

### Code availability

The code used to calculate NEG and replicate the analysis workflow is available at https://github.com/MZelko82/NEG.

### Author Contributions

**MZ:** conceptualisation, methodology, data curation, formal analysis, writing – original draft, review, and editing. **SR:** conceptualisation, methodology, writing – review and editing. **EH-Y:** conceptualisation, methodology, writing – review and editing. **HN:** conceptualisation, methodology, writing – review and editing.

