## Supplementary Materials for "Novel Exploration Growth Quantifies Anxiety-like Behaviour in the Elevated Plus Maze"

### Measuring Novel Exploration Phase Dynamics in the Elevated Plus Maze

#### Supplementary Materials:

#### S.1 EPM Grid Representation

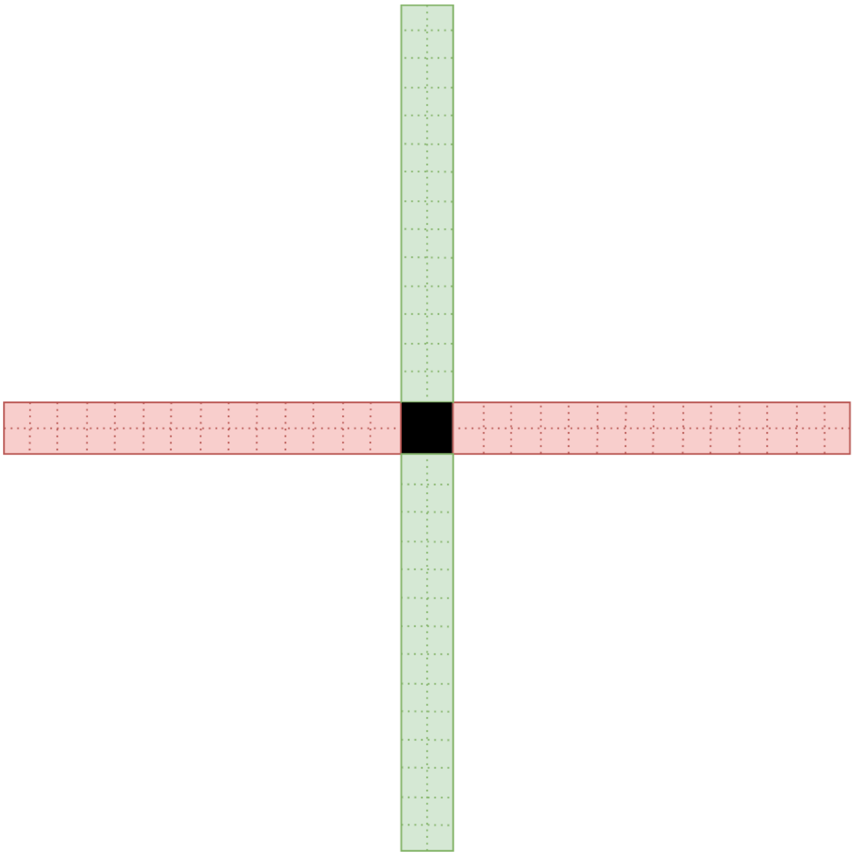

**Figure S-1** A Discretised Grid representing Exploration Visitation Points. The area of the maze is discretised into 1 $cm^2$ squares with the open arms are indicated in green, the closed arms in red. Note: grid is illustrative and not to scale.

#### S.2 Change-point Regression Model Specification

$$
\begin{aligned}
\mathrm{NEG}_i &\sim \mathrm{Binomial}(n_i, p_i) \\
\mathrm{logit}(p_i) &= \begin{cases} \beta_0 + \beta_1 \mathrm{time}_i, & 1 \le i \le k_1, \\ \alpha_0, & k_1 < i \le 300, \end{cases} \\
\beta_0 &\sim \mathrm{Normal}(-3, 3) \\
\beta_1 &\sim \mathrm{Beta}(1, 15) \\
\alpha_0 &\sim \mathrm{Normal}(0, 3) \\
k &\sim \mathrm{Dirichlet}(2)
\end{aligned}
\tag{1}
$$

$$
\begin{aligned}
\mathrm{NEG}_i &\sim \mathrm{Binomial}(n_i, p_i) \\
\mathrm{logit}(p_i) &= \begin{cases} \beta_0 + \beta_1 \mathrm{time}_i, & 1 \le i \le k_1, \\ \alpha_0, & k_1 < i \le k_2, \\ \gamma_0 + \gamma_1 \mathrm{time}_i, & k_2 < i \le 300, \end{cases} \\
\beta_0 &\sim \mathrm{Normal}(-3, 3) \\
\beta_1 &\sim \mathrm{Beta}(1, 15) \\
\alpha_0 &\sim \mathrm{Normal}(0, 3) \\
\gamma_0 &\sim \mathrm{Normal}(0, 3), \\
\gamma_1 &\sim \mathrm{Beta}(1, 15), \\
k &\sim \mathrm{Dirichlet}(2)
\end{aligned}
\tag{2}
$$

**Figure S-2** Model Specification for Single- and Dual-Phase Regression Models. (1) Single phase, and (2) dual phase exploration change-point regression models with prior distributions. $n_i = 50$, $k_i$ = change points.

#### S.3 Simulation Details for Novel Exploration Growth including Function Specifications

A single phase of bounded growth can be expressed using a logistic sigmoid function is expressed as:

$$y = \frac{f}{1+e^{(-ax+b)}} \tag{1}$$

where $f$ is a positive real integer, the domain $x \in (-\infty,\infty)$ the range is $[0, f]$. The growth of the curve is given by $a$ and the x-intercept of the inflexion point is located at $b$. This curve is characterised by a single phase of growth that increases toward an asymptote prior to or at the upper bound of the range. Conversely, a dual-phase growth curve can be expressed as:

$$y = f_{min} + \frac{f_{mid}-f_{min}}{1+e^{(-ax+b)}} + \frac{f_{max}-f_{mid}}{1+e^{(-cx+d)}} \tag{2}$$

where $f_{\text{min}}$, $f_{\text{mid}}$ and $f_{\text{max}}$ are positive real numbers where $f_{\text{min}} < f_{\text{mid}} < f_{\text{max}}$. This curve produces a similar initial growth phase which plateaus prior to the upper bound of the range. Following this plateau, a second growth phase occurs which grows toward an asymptote prior to or at the upper bound of the range.

$$\text{NEG}_T = \frac{NEG_{mid}}{1+e^{(-a \cdot time+b)}} + \frac{NEG_{max}}{1+e^{(-c \cdot time+d)}}$$

$$NEG_{mid,\ max} = 50,$$

$$a \sim Normal(0.2, 0.015), \qquad \text{Closed Arm Exploration Growth Rate}$$

$$b \sim Normal(5, 0.25), \qquad \text{Closed Arm Exploration Inflexion Time}$$

$$\left.\begin{array}{ll} c_E & \sim Normal(0.2, 0.015), \\ c_A & \sim Normal(1 \times 10^{-9}, 0), \\ c_D & \sim Normal(0.2, 0.001), \end{array}\right\} \rightarrow \qquad \text{Open Arm Exploration Growth Rate}$$

$$\left.\begin{array}{ll} d_E & \sim Normal(5, 0.25), \\ d_A & \sim Normal(100, 0), \\ d_D & \sim Normal(35, 0.25), \end{array}\right\} \rightarrow \qquad \text{Open Arm Exploration Inflexion Time}$$

**Figure S-3** Double Sigmoid Growth Model Specification and Simulation Distributions. The function $NEG_T$ specifies the exploration for the entire maze, with the distribution specifications for the simulation values for parameters *a, b, c,* and *d* for each phenotype. These parameter values were chosen as they produce reasonable exemplars for the expected time series for each phenotype. For example, the Exploratory Phenotype (E) shows rapid, arm-agnostic exploration growth at test onset, thus *a* and *c* are sampled from the same distribution, as are *b* and *d*. Conversely, the Delayed phenotype (D) has the same growth rate between the two arms; however, the inflexion time is different ($b < d_D$).

#### S.4 Simulation Details for Conventional Metrics

| Phenotype | Time in Open Arm *(s)* | Latency to Enter Open Arms *(ms)* | Entries into Open Arms *(n)* |
|---|---|---|---|
| Avoidant | 0 | 300 | 0 |
| Delayed | 100 | 185 | 2 |
| Exploratory | 100 | 5 | 2 |

$$y_i \sim \text{TruncNorm}(\mu = \mu_i,\ \sigma = \sigma_i), \qquad 0 < y_i < 300$$

**Figure S-4** Data Generating Process for Conventional Open Arm Measures. Values, y, for each measure were generated using a truncated normal distribution with bounds of 0 and 300. $\sigma_i = \mu_i \times 0.05$.

#### S.5 Conventional Metric Regression Summary

| | | 100% HDI | | | Region of Practical Equivalence (ROPE) | | |
|---|---|---|---|---|---|---|---|
| | Median | Low | High | PD | Region | HDI % | Decision |
| Latency to Enter Open Arms (s) | | | | | | | |
| Avoidant - Delayed | 98.03 | 99.13 | 96.92 | 100% | 11.78 | 0% | Rejected |
| Avoidant - Exploratory | 283.54 | 284.57 | 282.49 | 100% | 11.78 | 0% | Rejected |
| Delayed - Exploratory | 185.51 | 186.58 | 184.29 | 100% | 11.78 | 0% | Rejected |
| Open Arm Entries (n) | | | | | | | |
| Avoidant - Delayed | −2.00 | −1.99 | −2.01 | 100% | 0.09 | 0% | Rejected |
| Avoidant - Exploratory | −2.00 | −1.99 | −2.01 | 100% | 0.09 | 0% | Rejected |
| Delayed - Exploratory | 0.00 | 0.01 | −0.01 | 72% | 0.09 | 100% | Confirmed |
| Time Spent in Open Arms (s) | | | | | | | |
| Avoidant - Delayed | −100.07 | −99.51 | −100.66 | 100% | 4.73 | 0% | Rejected |
| Avoidant - Exploratory | −100.05 | −99.49 | −100.63 | 100% | 4.73 | 0% | Rejected |
| Delayed - Exploratory | 0.02 | 0.59 | −0.58 | 54% | 4.73 | 100% | Confirmed |

**Figure S-5** Regression Summary Table for Conventional Open Arm Measures. Equivalence is "confirmed" when the full posterior of the effect remains within the region, and "rejected" when less than 5% remains within the region. PD = Probability of Direction; HDI = Highest Density Interval;

### S.6 Prior Predictive Check for Single Phase Growth

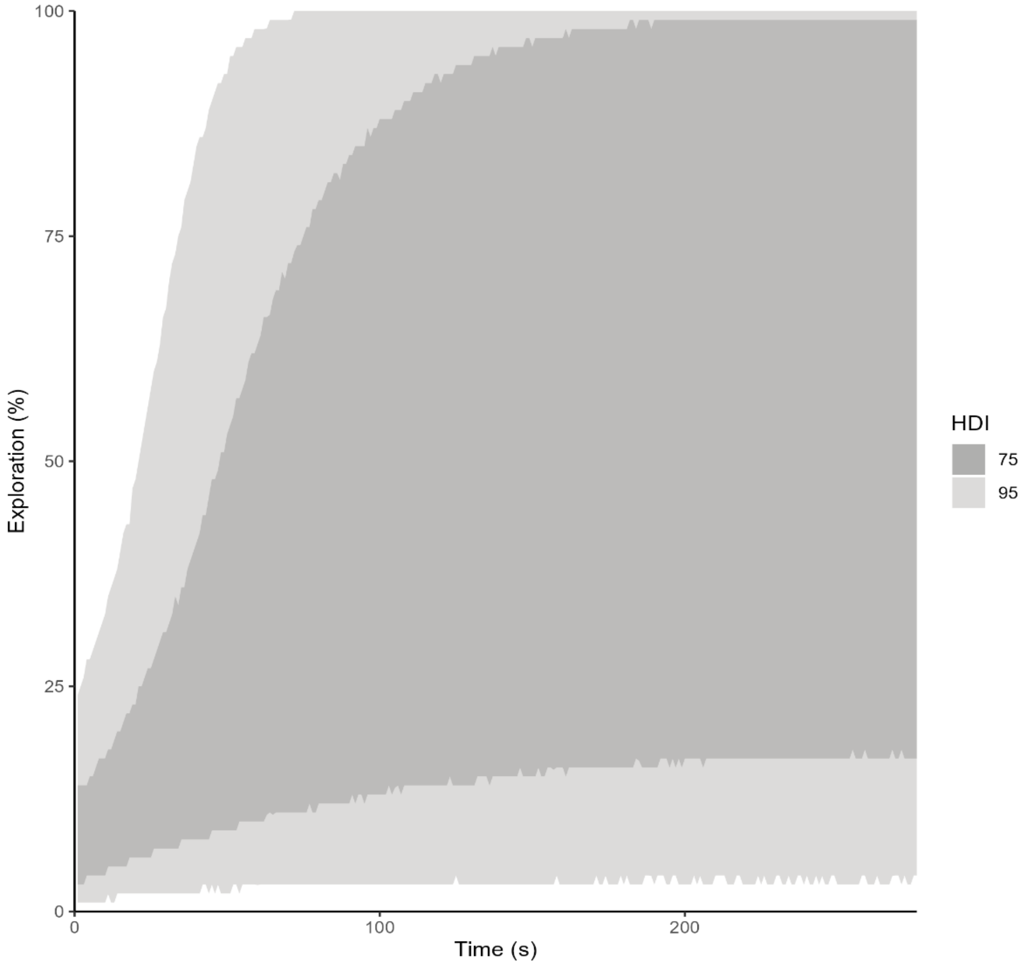


**Figure S-6** Prior Predictive Check: Single Phase Novel Exploration. Prior predictive checks demonstrate a suitable range of model outcomes given stated priors in model specifications. This check confirms that all time series trends are within the expected 0 to 100 % range.

#### S.7 Prior Predictive Check for Double Phase Growth

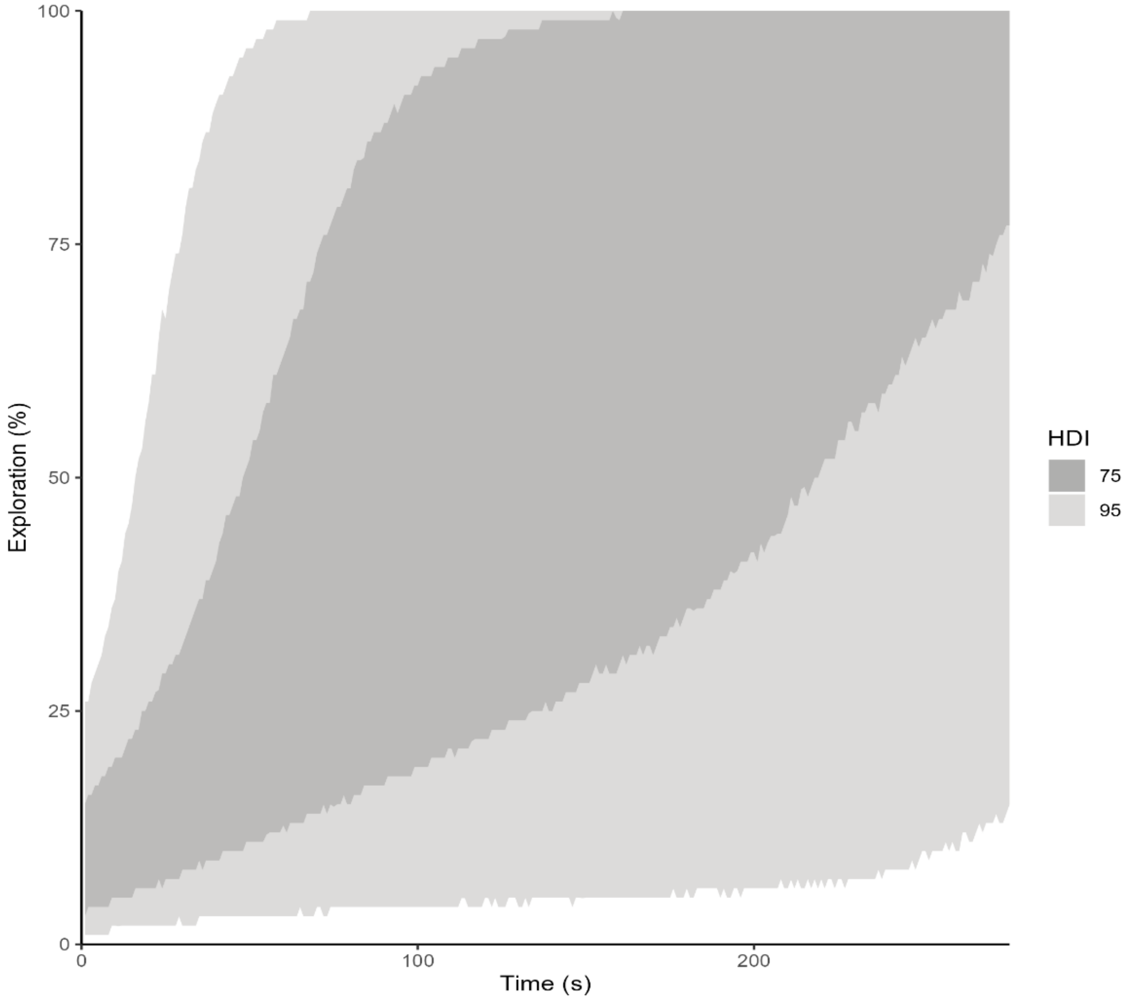


**Figure S-7** Prior Predictive Check: Double Phase Novel Exploration. Prior predictive checks demonstrate a suitable range of model outcomes given stated priors in model specifications. This check confirms that all time series trends are within the expected 0 to 100 % range.

### S.8 Change-point Regression Model Posterior Distributions and Diagnostics

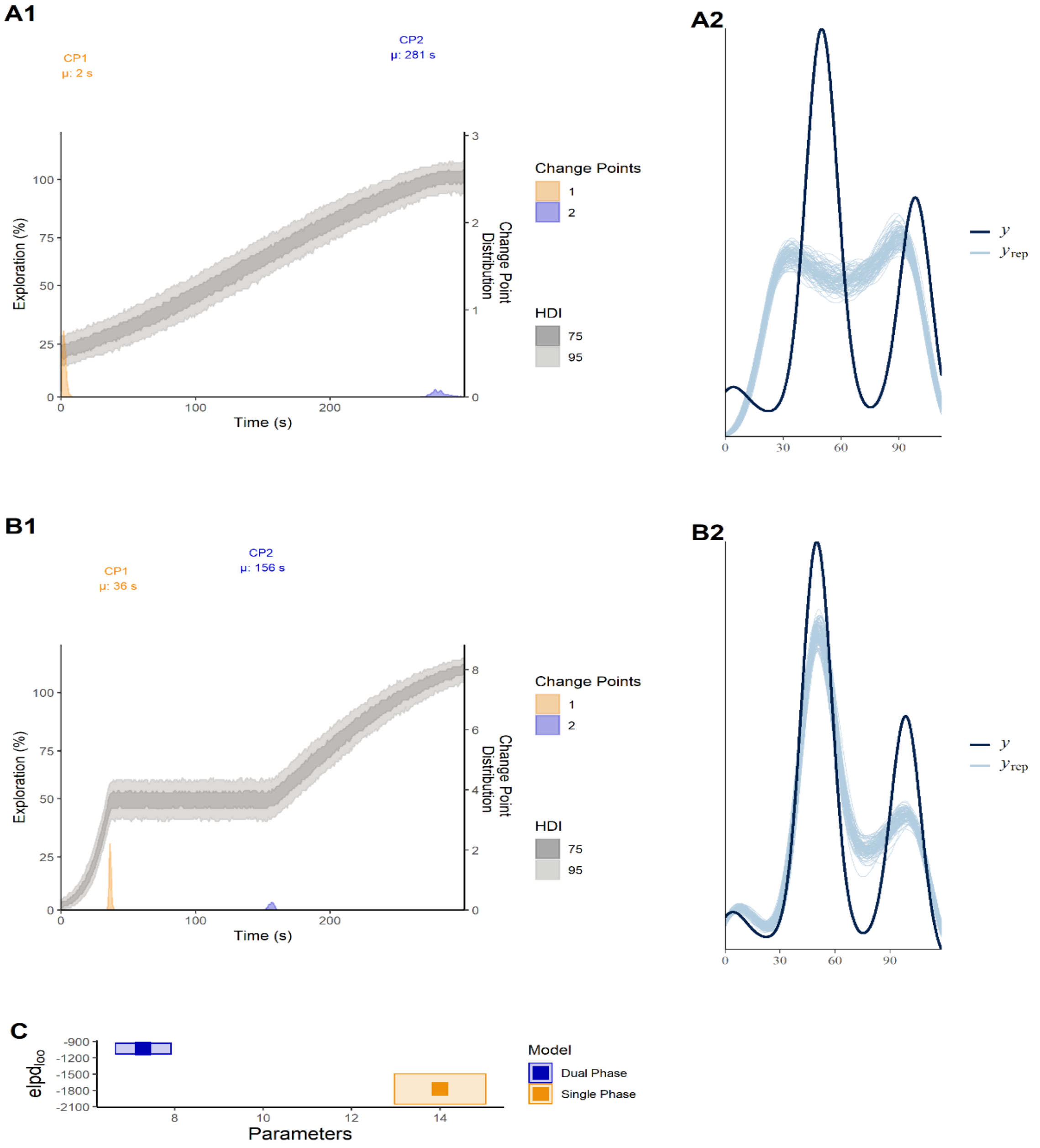

**Figure S-8a** Posterior Distributions for Single and Dual Phase Models of Total Exploration Growth by the Delayed Phenotype including Diagnostics. 75% and 95% posterior distributions for both single (A1) and dual phase models (B1) are displayed to evaluate the predicted trend and change points. Notably, the models disagree about the location of the change points. The dual phase model has greater predictive accuracy according to the posterior predictive checks (A2, B2); and, given the lower number of parameters and greater expected log pointwise predictive density ($elpd_{loo}$) via the LOOPS-CV comparison (C), it was preferred. CP1 = change point 1; CP2 = change point 1.

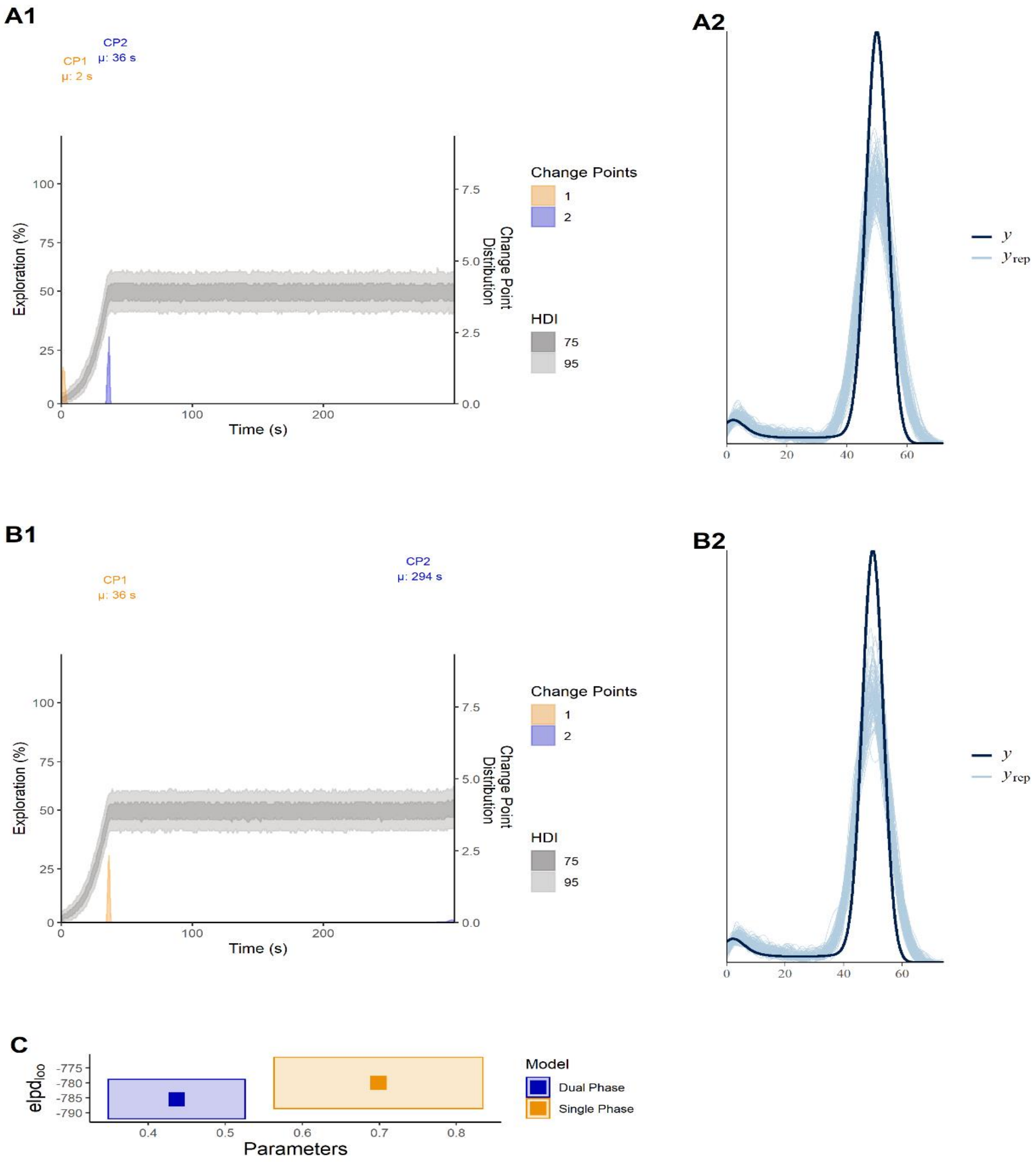
A1
CP1
μ: 2 s
CP2
μ: 36 s
Exploration (%)
Time (s)
Change Point Distribution
Change Points
1
2
HDI
75
95
A2
y
y_rep
B1
CP1
μ: 36 s
CP2
μ: 294 s
Exploration (%)
Time (s)
Change Point Distribution
Change Points
1
2
HDI
75
95
B2
y
y_rep
C
elpd_loo
Parameters
Model
Dual Phase
Single Phase

**Figure S-8b** Posterior Distributions of and Predictive Checks for Single and Dual Phase Change Point Modelling for Avoidant Phenotype Total Novel Exploration with Cross Validation Comparison. 75% and 95% posterior distributions for both single (A1) and dual phase models (B1) are displayed to evaluate the predicted trend and change points. Notably, the models agree about the location of the change points and share similar predictive accuracy according to the posterior predictive checks (A2, B2); thus, given the similar number of parameters and expected log pointwise predictive density ($elpd_{loo}$) via the LOOPS-CV comparison (C), the single phase model was preferred due to parsimony. CP1 = change point 1; CP2 = change point 1.

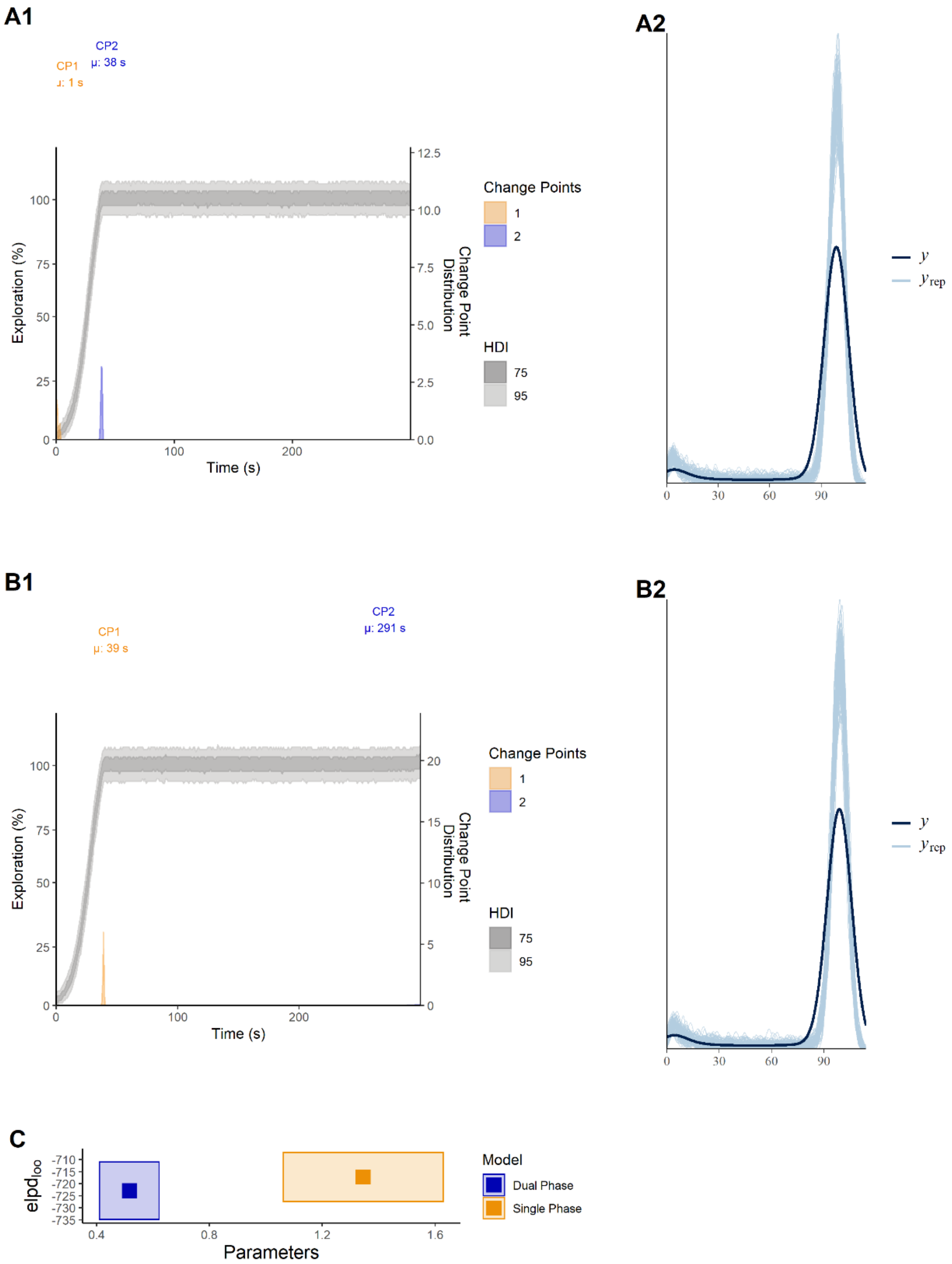
A1
CP2
μ: 38 s
CP1
μ: 1 s
Exploration (%)
Time (s)
Change Point Distribution
Change Points
1
2
HDI
75
95
A2
y
y_rep
B1
CP1
μ: 39 s
CP2
μ: 291 s
Exploration (%)
Time (s)
Change Point Distribution
Change Points
1
2
HDI
75
95
B2
y
y_rep
C
elpd_loo
Parameters
Model
Dual Phase
Single Phase

**Figure S-8c** Posterior Distributions of and Predictive Checks for Single and Dual Phase Change Point Modelling for Exploratory Phenotype Total Novel Exploration with Cross Validation Comparison. 75% and 95% posterior distributions for both single (A1) and dual phase models (B1) are displayed to evaluate the predicted trend and change points. Notably, the models agree about the location of the change points and share similar predictive accuracy according to the posterior predictive checks (A2, B2); thus, given the similar number of parameters and expected log pointwise predictive density ($elpd_{loo}$) via the LOOPS-CV comparison (C), the single phase model was preferred due to parsimony. CP1 = change point 1; CP2 = change point 1.

#### S.9 Total Maze Exploration General Additive Regression Model Specification

$$
\begin{aligned}
NEG_i &\sim \text{Normal}(\mu_i, \sigma) \\
\mu_i &= f(x_i) \\
f(x_i) &= \beta_1 \text{group}_i + s_j \text{time}_i + \alpha_{n_i}, \qquad \text{for } 1,\ldots,\ J \text{ basis terms } (time) \\
\beta &\sim Normal(0,1) \\
s_j &\sim Normal(0, \sigma_\tau) \\
\alpha_n &\sim \text{Normal}(\alpha, 1) \\
\alpha &\sim \text{Normal}(0,1) \\
\sigma &\sim \text{Exponential}(1) \\
\sigma_\tau &\sim \text{Exponential}(1)
\end{aligned}
$$

**Figure S-9** Model Specification for General Additive Regression of Total Exploration by Phenotype including Priors. $j = 30$.

### S.10 Posterior Predictive Check for Total Maze Exploration Regression Model

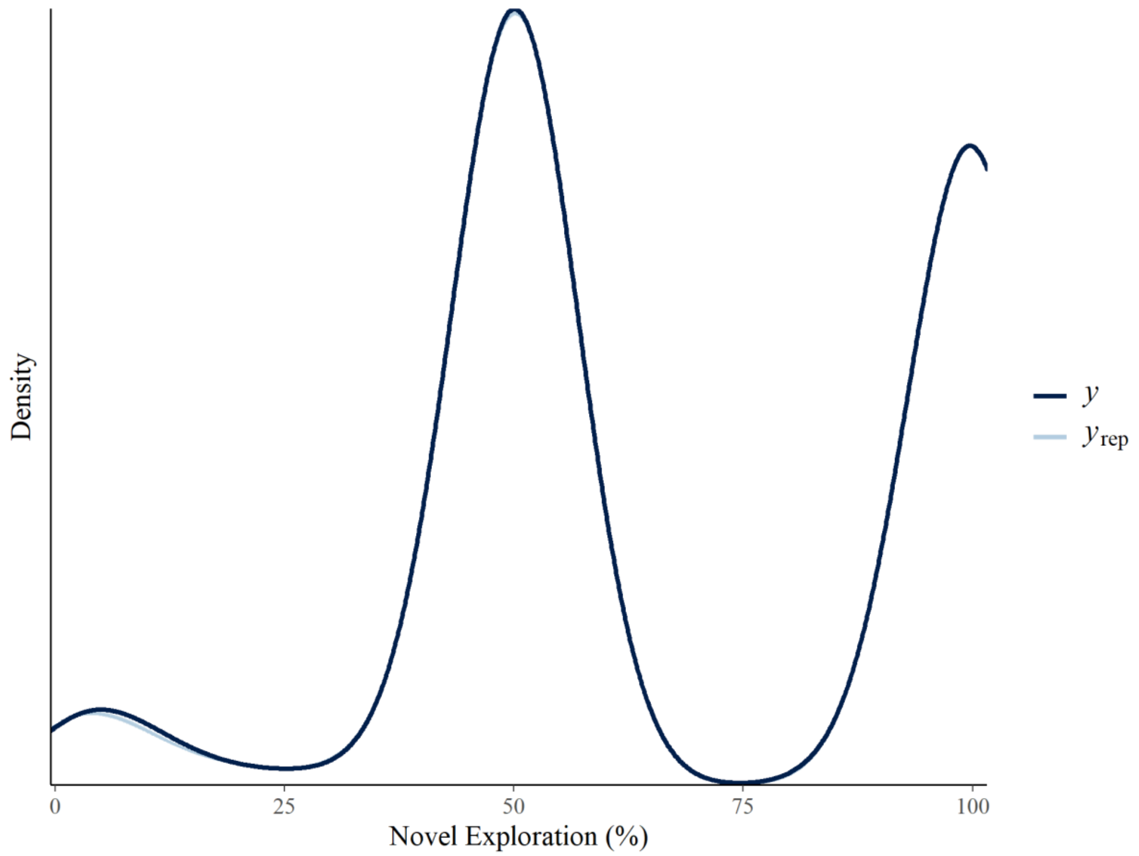


**Figure S-10** Posterior Predictive Check for Non-Linear Regression of Total Exploration Growth by Phenotype. $y$ represents the data, and $y_{rep}$ represents the predicted outcome based on the model fit.

#### S.11 Total Maze Exploration General Additive Regression Summary

| | | | 95% HDI | | | |
|---|---|---|---|---|---|---|
| Phenotype | Median | MAD | High | Low | Rhat | Bulk ESS |
| *Group* | | | | | | |
| Avoidant | −0.71 | 0.00 | −0.71 | −0.70 | 1.00 | 5,185.83 |
| Delayed | −0.04 | 0.00 | −0.04 | −0.04 | 1.00 | 5,034.72 |
| Exploratory | 0.75 | 0.00 | 0.74 | 0.75 | 1.00 | 5,174.42 |
| *Smooth(Time)* | | | | | | |
| Avoidant | 0.21 | 0.14 | −0.06 | 0.48 | 1.00 | 8,487.11 |
| Delayed | 0.22 | 0.14 | −0.06 | 0.50 | 1.00 | 8,707.34 |
| Exploratory | 0.41 | 0.14 | 0.13 | 0.69 | 1.00 | 12,205.24 |

**Figure S-11** Regression Summary for Total Novel Exploration Growth by Phenotype.

Note: MAD = Median Absolute Deviation, HDI = Highest Density Interval, ESS = Effective Sample Size.

### S.12 Change-point Regression Posterior Distributions and Diagnostics

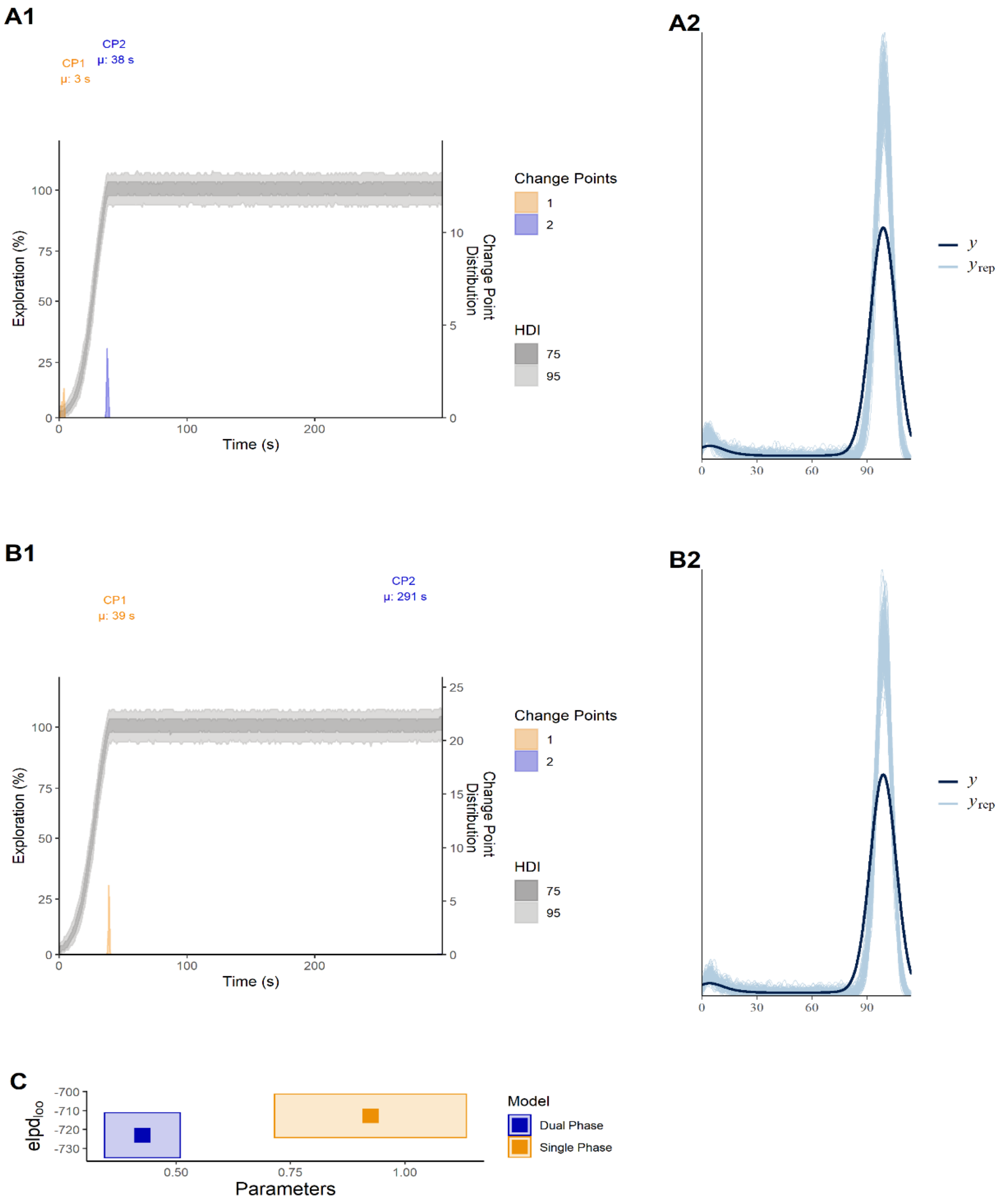

**Figure S-12a** Posterior Distributions of and Predictive Checks for Single and Dual Phase Change Point Modelling for Avoidant Phenotype Novel Exploration of Closed Arms with Cross Validation comparison. 75% and 95% posterior distributions for both single (A1) and dual phase models (B1) are displayed to evaluate the predicted trend and change points. Notably, the models agree about the location of the change points and share similar predictive accuracy according to the posterior predictive checks (A2, B2); thus, given the similar number of parameters and expected log pointwise predictive density ($elpd_{loo}$) via the LOOPS-CV comparison (C), the single phase model was preferred due to parsimony. CP1 = change point 1; CP2 = change point 1.

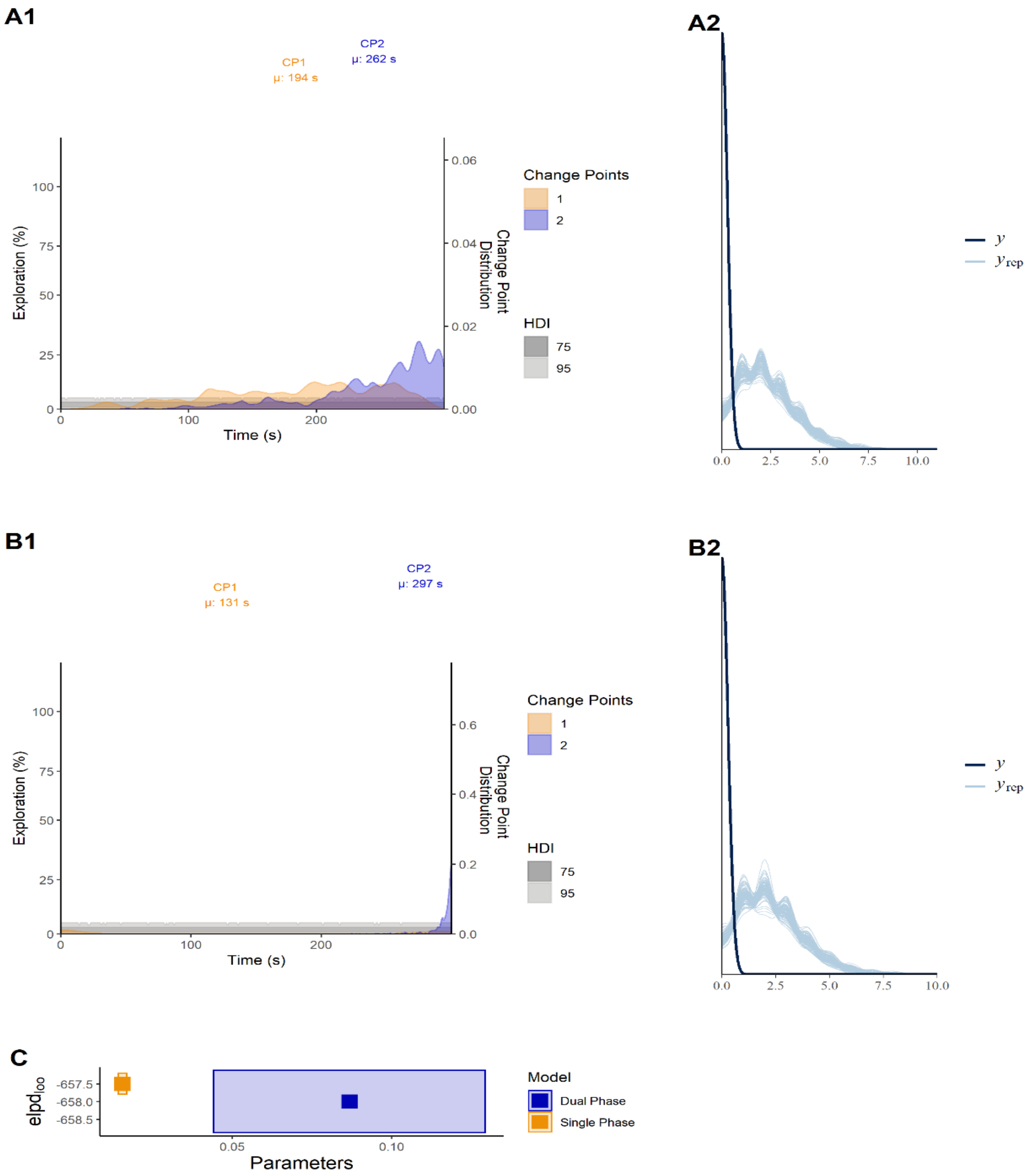
A1
CP1
μ: 194 s
CP2
μ: 262 s
Exploration (%)
Time (s)
Change Point Distribution
Change Points
1
2
HDI
75
95
A2
y
y_rep
B1
CP1
μ: 131 s
CP2
μ: 297 s
Exploration (%)
Time (s)
Change Point Distribution
Change Points
1
2
HDI
75
95
B2
y
y_rep
C
elpd_loo
Parameters
Model
Dual Phase
Single Phase

**Figure S-12b** Posterior Distributions of and Predictive Checks for Single and Dual Phase Change Point Modelling for Avoidant Phenotype Novel Exploration of Open Arms with Cross Validation comparison. 75% and 95% posterior distributions for both single (A1) and dual phase models (B1) are displayed to evaluate the predicted trend and change points. Notably, neither model locates of the change points accurately and share similar predictive accuracy according to the posterior predictive checks (A2, B2); thus, given the similar number of parameters and expected log pointwise predictive density ($elpd_{loo}$) via the LOOPS-CV comparison (C), the single phase model was preferred due to parsimony. CP1 = change point 1; CP2 = change point 1.

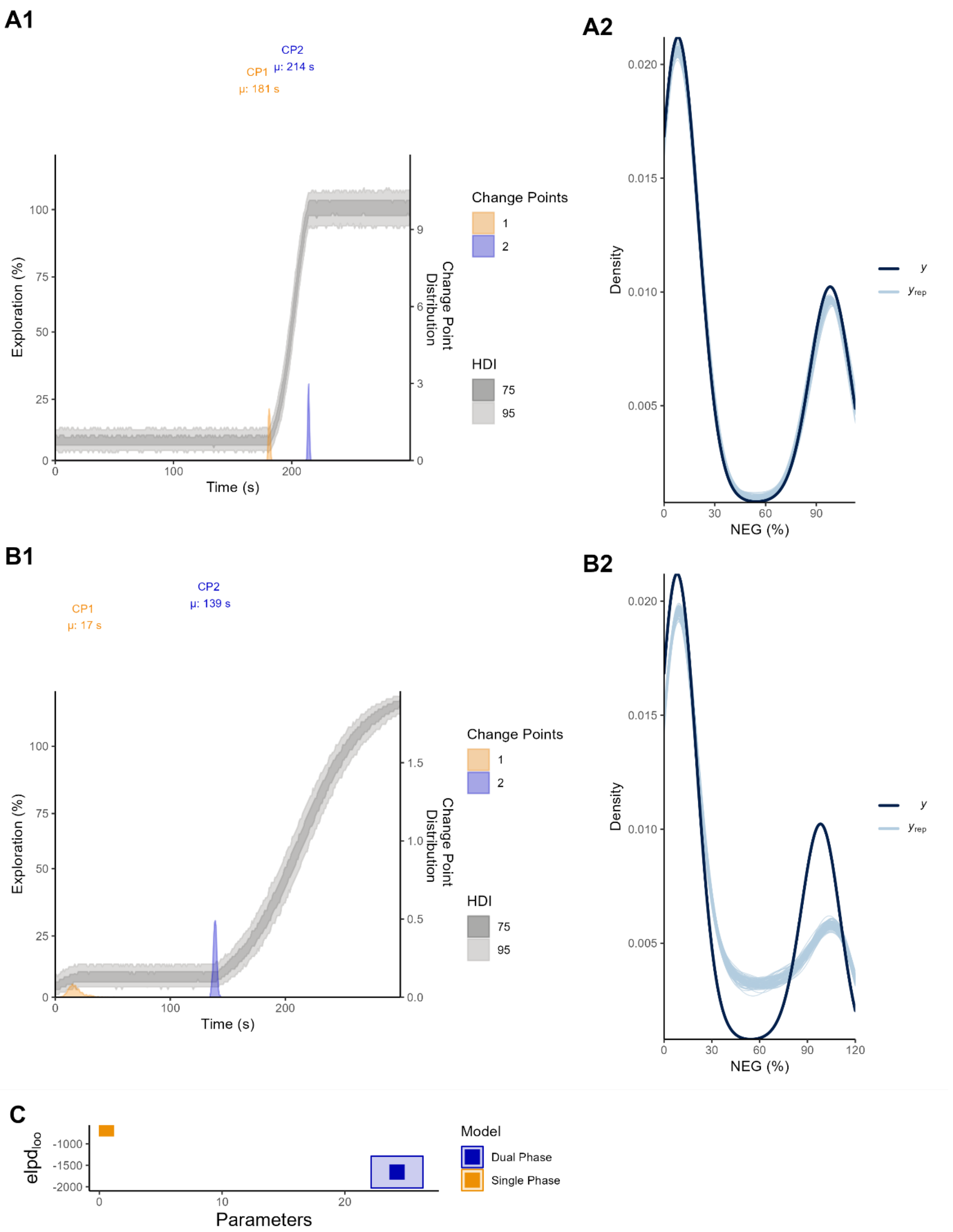
A1
CP2
μ: 214 s
CP1
μ: 181 s
Exploration (%)
Time (s)
Change Point Distribution
Change Points
1
2
HDI
75
95
A2
Density
NEG (%)
y
y_rep
B1
CP1
μ: 17 s
CP2
μ: 139 s
Exploration (%)
Time (s)
Change Point Distribution
Change Points
1
2
HDI
75
95
B2
Density
NEG (%)
y
y_rep
C
elpd_loo
Parameters
Model
Dual Phase
Single Phase

**Figure S-12c** Posterior Distributions of and Predictive Checks for Single and Dual Phase Change Point Modelling for Delayed Phenotype Open Arm Novel Exploration with Cross Validation Comparison. 75% and 95% posterior distributions for both single (A1) and dual phase models (B1) are displayed to evaluate the predicted trend and change points. Notably, the models disagree about the location of the change points, with the single phase reporting higher predictive accuracy according to the posterior predictive checks (A2, B2); thus, given the smaller number of parameters and greater expected log pointwise predictive density ($elpd_{loo}$) via the LOOPS-CV comparison (C), the single phase model was preferred. CP1 = change point 1; CP2 = change point 1.

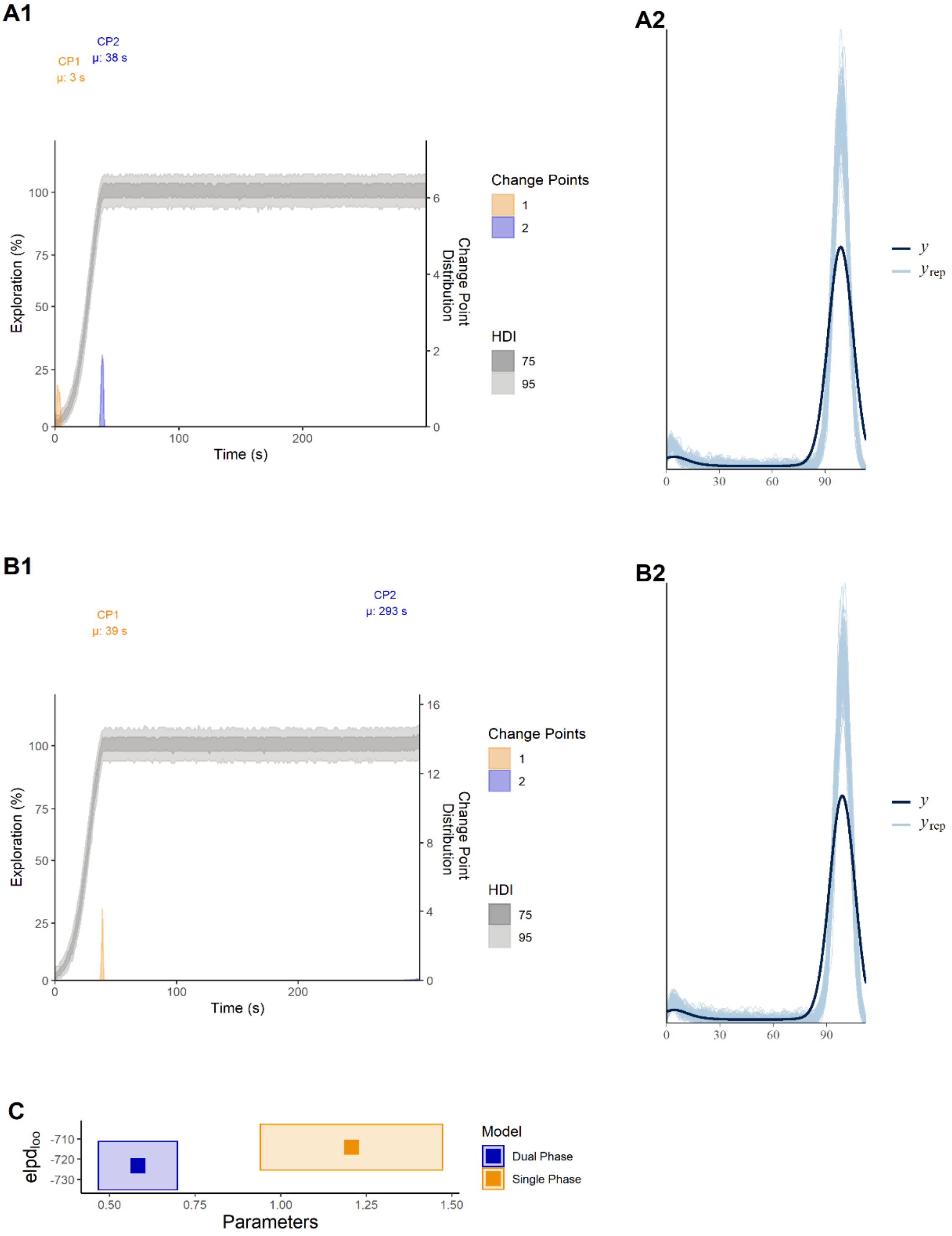
A1
CP1
μ: 3 s
CP2
μ: 38 s
Exploration (%)
Time (s)
Change Point Distribution
Change Points
1
2
HDI
75
95
A2
y
y_rep
B1
CP1
μ: 39 s
CP2
μ: 293 s
Exploration (%)
Time (s)
Change Point Distribution
Change Points
1
2
HDI
75
95
B2
y
y_rep
C
elpd_loo
Parameters
Model
Dual Phase
Single Phase

**Figure S-12d** Posterior Distributions of and Predictive Checks for Single and Dual Phase Change Point Modelling for Delayed Phenotype Novel Exploration of Closed Arms with Cross Validation Comparison. 75% and 95% posterior distributions for both single (A1) and dual phase models (B1) are displayed to evaluate the predicted trend and change points. Notably, the models agree about the location of the change points and share similar predictive accuracy according to the posterior predictive checks (A2, B2); thus, given the similar number of parameters and expected log pointwise predictive density ($elpd_{loo}$) via the LOOPS-CV comparison (C), the single phase model was preferred due to parsimony. CP1 = change point 1; CP2 = change point 1.

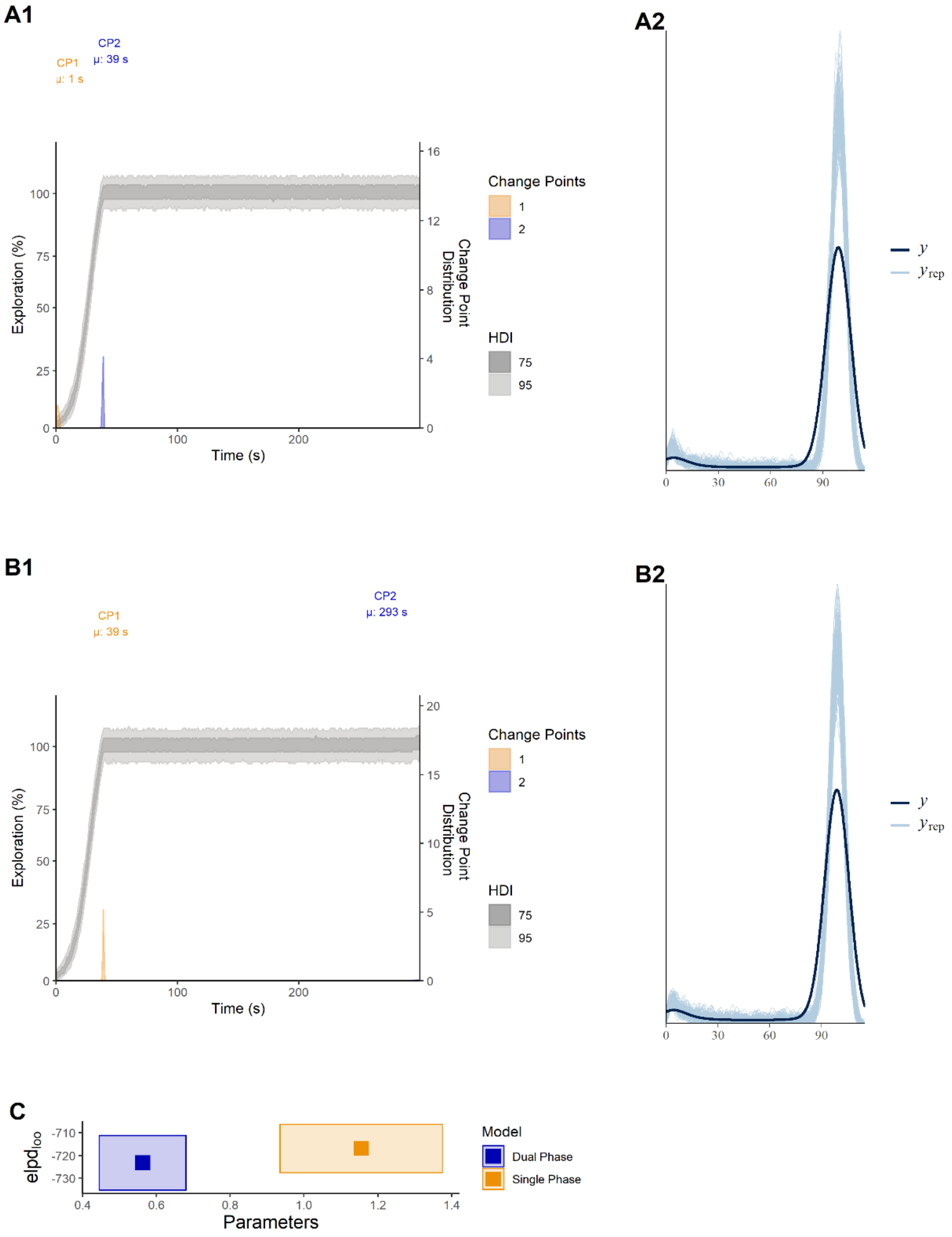
A1
CP1
µ: 1 s
CP2
µ: 39 s
Exploration (%)
Time (s)
Change Point Distribution
Change Points
1
2
HDI
75
95
A2
y
y_rep
B1
CP1
µ: 39 s
CP2
µ: 293 s
Exploration (%)
Time (s)
Change Point Distribution
Change Points
1
2
HDI
75
95
B2
y
y_rep
C
elpd_loo
Parameters
Model
Dual Phase
Single Phase

**Figure S-12e** Posterior Distributions of and Predictive Checks for Single and Dual Phase Change Point Modelling for Exploratory Phenotype Novel Exploration of Closed Arms with Cross Validation Comparison. 75% and 95% posterior distributions for both single (A1) and dual phase models (B1) are displayed to evaluate the predicted trend and change points. Notably, the models agree about the location of the change points and share similar predictive accuracy according to the posterior predictive checks (A2, B2); thus, given the similar number of parameters and expected log pointwise predictive density (elpdloo) via the LOOPS-CV comparison (C), the single phase model was preferred due to parsimony. CP1 = change point 1; CP2 = change point 1.

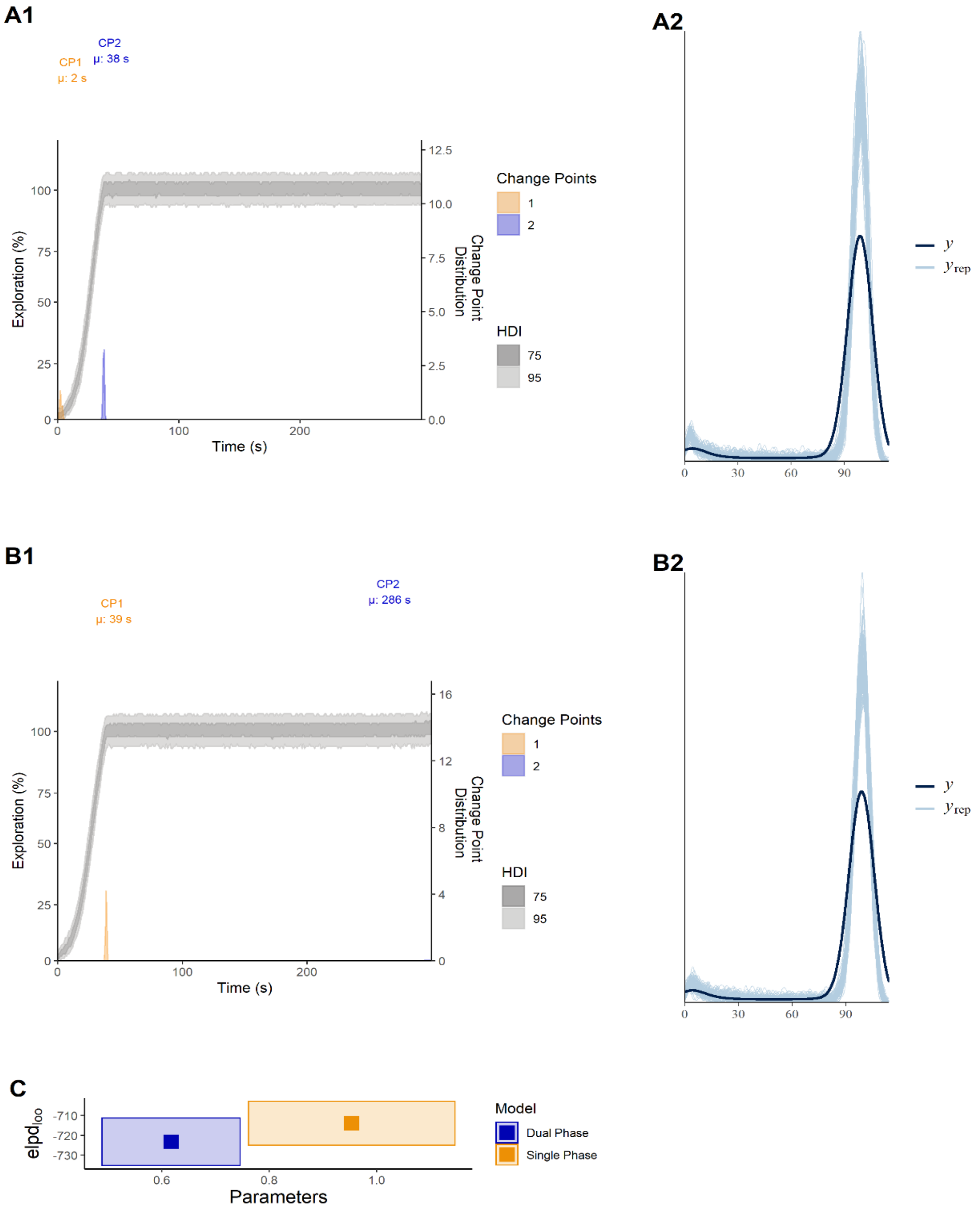
A1
CP1
μ: 2 s
CP2
μ: 38 s
Exploration (%)
Time (s)
Change Point Distribution
Change Points
1
2
HDI
75
95
A2
y
$y_{rep}$
B1
CP1
μ: 39 s
CP2
μ: 286 s
Exploration (%)
Time (s)
Change Point Distribution
Change Points
1
2
HDI
75
95
B2
y
$y_{rep}$
C
$elpd_{loo}$
Parameters
Model
Dual Phase
Single Phase

**Figure S-12f** Posterior Distributions of and Predictive Checks for Single and Dual Phase Change Point Modelling for Exploratory Phenotype Novel Exploration of Open Arms with Cross Validation Comparison. 75% and 95% posterior distributions for both single (A1) and dual phase models (B1) are displayed to evaluate the predicted trend and change points. Notably, the models agree about the location of the change points and share similar predictive accuracy according to the posterior predictive checks (A2, B2); thus, given the similar number of parameters and expected log pointwise predictive density (elpdloo) via the LOOPS-CV comparison (C), the single phase model was preferred due to parsimony. CP1 = change point 1; CP2 = change point 1.

#### S.13 Arm Exploration Regression Model Specification

$$\text{NEG}_i \sim \text{Binomial}(n_i, p_i)$$

$$\text{logit}(p_i) = \alpha_{j_i} + \beta_{j_i}\text{time}_i \times \beta_3\text{phenotype}_i \times \beta_2\text{arm}_i$$

$$\begin{bmatrix} \alpha_j \\ \beta_j \end{bmatrix} \sim \text{MVNormal}\left( \begin{bmatrix} \alpha \\ \beta \end{bmatrix}, \mathbf{S} \right)$$

$$\mathbf{S} = \begin{pmatrix} \sigma_\alpha & 0 \\ 0 & \sigma_\beta \end{pmatrix} \mathbf{R} \begin{pmatrix} \sigma_\alpha & 0 \\ 0 & \sigma_\beta \end{pmatrix}$$

$$\alpha,\ \beta \sim \text{Normal}(0, 1.5)$$

$$\sigma_\alpha,\ \sigma_\beta \sim \text{HalfCauchy}(0, 1)$$

$$\mathbf{R} \sim \text{LKJcorr}(2)$$

**Figure S-13** Model Specification for Generalised Linear Mixed Effects Regression of Novel Exploration by Arm per Phenotype including priors. $n = 50$, $i =$ value at timepoint, $j =$ subject.

#### S.14 Arm Exploration Regression Model Summary

| | | | 100% HDI | | | |
|---|---|---|---|---|---|---|
| Phenotype | Median | MAD | Low | High | Rhat | ESS |
| *Closed Arm* | | | | | | |
| Avoidant | -4.66 | 0.12 | -4.90 | -4.42 | 1.00 | 2588 |
| Delayed | 8.49 | 0.24 | 8.01 | 8.98 | 1.00 | 2827 |
| Exploratory | 0.82 | 0.23 | 0.37 | 1.27 | 1.00 | 2324 |
| *Time: Closed Arm* | | | | | | |
| Avoidant | 0.18 | 0.00 | 0.17 | 0.19 | 1.00 | 2339 |
| Delayed | -0.05 | 0.01 | -0.06 | -0.04 | 1.00 | 2132 |
| Exploratory | -0.18 | 0.01 | -0.20 | -0.17 | 1.00 | 2212 |
| *Open Arm* | | | | | | |
| Avoidant | -3.89 | 0.08 | -4.05 | -3.74 | 1.00 | 3281 |
| Delayed | -8.22 | 0.19 | -8.57 | -7.85 | 1.00 | 3339 |
| Exploratory | -1.00 | 0.15 | -1.28 | -0.70 | 1.00 | 2426 |
| *Time: Open Arm* | | | | | | |
| Avoidant | 0.00 | 0.00 | 0.00 | 0.00 | 1.00 | 3252 |
| Delayed | 0.06 | 0.00 | 0.06 | 0.06 | 1.00 | 3257 |
| Exploratory | 0.19 | 0.00 | 0.18 | 0.20 | 1.00 | 1981 |

**Figure S-14** Fixed Effects Summary for Novel Exploration Growth by Arm per Phenotype. Note: Values are reported on the response scale. MAD = Median Absolute Deviation, HDI = Highest Density Interval, ESS = Effective Sample Size.
